# An Anisotropic Reactive Viscoelastic Model of the Rhesus Macaque Cervix for Studying Cervical Remodeling

**DOI:** 10.1101/2025.05.14.654071

**Authors:** Camilo A. Duarte-Cordon, Shuyang Fang, Ivan M. Rosado-Mendez, Gerard Ateshian, Timothy J. Hall, Helen Feltovich, Kristin M. Myers

## Abstract

The uterine cervix is a soft biological tissue with critical biomechanical functions in pregnancy. It is a mechanical barrier that supports the growing fetus. As pregnancy progresses, the cervix becomes more compliant and eventually opens in late pregnancy to facilitate childbirth. This dual function is facilitated by extensive remodeling of the cervical extracellular matrix (ECM), giving rise to its complex time-dependent material properties. Premature cervical remodeling is known to result in preterm birth, defined as birth before 37 weeks of gestation. While previous work has studied cervical remodeling using various biomechanical methods, it remains unclear how the intrinsic or flow-independent viscoelastic behavior of the cervix is influenced by cervical remodeling. In this study, an anisotropic reactive viscoelastic material model was formulated and investigated under tensile deformation to understand material behavior in cervical remodeling. To calibrate the model, experimental force relaxation data was used from uniaxial tension tests on Rhesus macaque cervical specimens from four gestational time points. The results showed that cervical tissue equilibrium and instantaneous stiffness significantly decreased from the non-pregnant to the late pregnancy status. In addition, cervical tissue in the late third trimester relaxed faster to equilibrium than the other gestational groups, particularly at prescribed tensile strains greater than 30%. This fast relaxation to equilibrium helps the cervix dissipate tensile hoop stresses induced by the fetus during labor, preventing its rupture. This work provides insights into time-dependent cervical remodeling features, which are crucial for developing diagnostic methods and treatments for preterm birth.

## 1 Introduction

The uterine cervix is a soft biological tissue located at the lower end of the uterus that performs critical biomechanical functions in pregnancy[1] and fertility[2; 3]. During pregnancy, the cervix is a mechanical barrier, initially stiff and closed, that protects and keeps the growing fetus in the uterus. As labor approaches, the cervix shortens and becomes a more compliant structure, allowing it to dilate significantly for childbirth at term. This remarkable function of the cervix is facilitated by extensive remodeling, i.e., changes in the structure and composition throughout pregnancy, of its extracellular matrix (ECM) [4; 5]. The structure of the cervical ECM, mainly composed of a three-dimensional network of fibrillar collagen types I and III, glycosaminoglycans (GAGs), proteoglycans (PGs), elastin, spatially varying cellular content (e.g., smooth muscle cells (SMC)), and interstitial fluid [6; 7; 8], gives rise to its complex anisotropic and time-dependent material properties, including nonlinear elasticity [9; 10], intrinsic viscoelasticity (fluid flow independent) [11; 12], and poroelasticity [13]. Notably, changes in the tissue-level equilibrium stiffness and compressive poroviscoelasticity of the murine and human cervix have been associated with different stages of normal and disrupted cervical remodeling [14; 15; 16; 10; 17; 18; 13]. Nevertheless, it remains unclear how the intrinsic tensile viscoelastic behavior of the cervix is affected in pregnancy. Gaining insights into the changes in cervical intrinsic viscoelasticity throughout gestation could not only deepen our knowledge of cervical remodeling but also help identify biomarkers for diagnosing cervical pathophysiologies involving inflammation and premature softening [19], such as cervical insufficiency, associated with an increased risk of spontaneous preterm birth (sPTB) (birth before 37 weeks of gestation) [20].

Similar to other soft biological tissues [21], the cervix is viscoelastic, as it exhibits time-dependent mechanical behavior, which can be observed in stress-relaxation and creep experiments. [12; 11; 13; 22]. Depending on the molecular mechanism and type of stress applied (e.g., compression, tension, and/or shear), cervical tissue can exhibit intrinsic or flow-independent viscoelasticity, poroelasticity, or a combination of these. Intrinsic viscoelasticity in soft collagenous materials such as the cervix is associated with molecular mechanisms such as crimping/uncrimping of collagen fibers [23], and intermittent breaking of weak inter-/intra-molecular bonds or entanglements within the collagen fiber network [24; 25]. In contrast, poroelasticity involves frictional interactions between the interstitial fluid and solid components of the ECM [26; 27]. Under uniaxial tensile loading, the mechanical response of soft tissues is expected to be dominated by the collagen fiber network as it bears most of the load, and therefore, poroelasticity is less critical. Quantifying changes in the intrinsic viscoelastic properties of the cervix during gestation could enhance our understanding of the role of the collagen network structure in cervical remodeling.

Engineering tools, such as constitutive modeling and mechanical testing of the cervix, have made it possible to obtain valuable insights into cervical tissue remodeling in terms of alterations in its tissue-level material properties. Hyperelastic material models and inverse finite element analysis (IFEA) frame-works, incorporating mechanical testing data from murine and human cervical specimens, have allowed researchers to model the nonlinear elasticity of cervical tissue and quantify changes in its equilibrium material properties throughout pregnancy [14; 28; 15; 16; 10; 17; 29]. Furthermore, changes in the steady-state material properties of the cervix, such as cervical collagen fiber stiffness, have been correlated with alterations in the structure of the cervical ECM [14; 28; 19]. For instance, studies using *ex vivo* cervical tissue from mice have shown that equilibrium collagen fiber and ground substance stiffness decrease significantly after the 6th day of gestation [28]. This drop in mouse cervical stiffness through pregnancy has been associated with a decrease in intermolecular collagen crosslinks maturity ratio (ratio of mature to immature crosslinks) [14; 30], an increase in disorganization in the collagen network [31], and a drastic increase in hyaluronic acid (HA), a type of GAG, particularly in late pregnancy [19; 32]. Additionally, the lack of PGs, such as decorin and biglycan, in cervical tissue has been shown to reduce the extensibility of the cervix, highlighting their critical role in cervical remodeling [16].

Previous studies on stress relaxation and creep in cervical tissue indicate that the viscoelasticity of the cervix also changes during pregnancy [22; 33; 34; 35; 36; 37; 38; 17; 33]. Harkness and Harkness [22] found that near-term rat cervices were more extensible and crept more under constant tensile loading than non-pregnant cervices. In stress relaxation tests performed on the cervical canal of patients with and without cervical insufficiency (CI) using a water-filled balloon, van Duyl et al. [33] observed that patients with CI had significantly longer stress relaxation times than healthy patients. Barone et al. [34] found that late-pregnant rat cervical tissue relaxed faster to equilibrium under unconfined compression than non-pregnant or mid-pregnant specimens. Ashofteh et al. [36] reported similar findings in force relaxation tests under uniaxial tension, observing that equivalent to ”third-trimester” rat cervical tissue relaxed faster than non-pregnant tissue. Yao et al. [35] found that under spherical indentation, the equilibrium and instantaneous stiffness of human cervical tissue specimens were lower in pregnant tissue specimens than in non-pregnant ones. Shi and Myers [13] employed a poro-viscoelastic model to analyze stress relaxation from spherical indentation, revealing that non-pregnant specimens had much lower hydraulic permeability than pregnant specimens, with the external os exhibiting higher permeability than the internal os. Various rheological models, including the Maxwell, Kelvin–Voigt, and Zener models, have been used to study the cervix’s viscoelastic properties and its frequency dependence [37; 38; 39]. However, most of these models treat the cervix as a homogeneous and linearly elastic material and do not provide much information on its anisotropic viscoelastic properties. This highlights the need for methods that can effectively characterize the cervix’s anisotropic viscoelasticity.

Although previous studies have shed light on the changes in the time-dependent properties of the cervix across gestation, they have primarily focused on its compressive properties and utilized both murine and human cervical tissue, which have some limitations. Rodents, such as mice and rats, despite having differences in uterine anatomy compared to humans, are valuable animal models for studying cervical remodeling as they exhibit cervical ECM remodeling patterns that resemble those in humans and allow the analysis of multiple mid-gestational time points [40; 30]. Studying cervical remodeling using *ex vivo* human cervical tissue remains challenging, and it is usually limited to two gestational points (usually term pregnant and non-pregnant). Non-human primate models, particularly the Rhesus macaque, are similar to humans in terms of maternal-fetal anatomy and physiology [41; 42], and they are well-established animal models that have been used to study infection-induced preterm labor [43]. Furthermore, the *in vivo* cervical stiffness in Rhesus macaques has been measured using quantitative ultrasound parameters (QUS), such as shear wave speed (SWS), obtained from shear wave elastography imaging (SWEI) [18; 44]. The stiffness of soft biological tissues can be inferred from SWS, as shear waves propagate faster in stiffer tissue. SWE studies on Rhesus indicate that SWS-derived cervical stiffness during late pregnancy decreases to less than 40 % of pre-pregnancy levels and recovers two weeks after delivery [18]. The stiffness calculated using SWS often relies on assumptions of tissue linear elasticity and homogeneity, which do not apply to the cervix, as it is a highly anisotropic and viscoelastic biological tissue. However, accurate SWS measurement, appropriate assumptions of boundary conditions, and accounting for the effect of cervical tissue heterogeneity and viscosity can help us provide reliable estimates of cervical tissue stiffness [44] and evaluate the potential of SWEI-derived parameters, such as SWS, as biomarkers of cervical remodeling.

This study aims to understand how the time-dependent material properties of the Rhesus macaque cervix, subjected to *ex vivo* uniaxial tension, evolve with cervical remodeling at four relevant gestational time points. To accomplish this goal, a three-dimensional anisotropic reactive viscoelastic model [45] of the Rhesus macaque cervix is implemented. The model was calibrated with force relaxation data from uniaxial tension and spherical indentation testing at three strains[10]. This paper is structured as follows: Section 2 details the constitutive model used to simulate the relaxation response of the Rhesus macaque cervical tissue, the inverse modeling approach used to obtain the material parameters of the model, the model validation approach, and the statistical methods used to assess significant differences in material properties. The numerical results of this study are presented in Section 3 and discussed in Section 4. Finally, Section 5 provides a summary and conclusions of this study.

## 2 Materials and Methods

### 2.1 Constitutive Material Model

Inspired by the solid components of the cervical ECM, cervical tissue from the Rhesus macaque was modeled as an anisotropic solid material with a viscoelastic collagen fiber network [28; 16; 45]. For this purpose, a constitutive material model was formulated for large deformation consisting of a solid mixture of a reactive viscoelastic continuous distributed collagen fiber network and a compressible neo-Hookean ground substance. The total strain energy density of the material is given by:

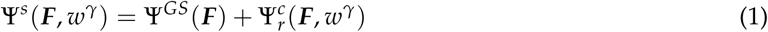

where Ψ*^GS^* and 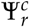 represent the contributions from the ground substance and collagen fiber network, ***F*** is the deformation gradient with Jacobian *J* = det ***F***. The ground substance was assumed to only contribute to the equilibrium elastic response of the tissue, and it represents the mechanical response of non-fibrillar ECM components, such as GAGs and PGs. The strain energy density of the ground substance is given by [46]:

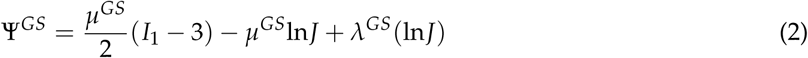

where *µ^GS^* and *λ^GS^* are the Lamé constants, which can be used to obtain the Young’s modulus *E^GS^* = *µ^GS^*(3*λ^GS^* + 2*µ^GS^*)/(*λ^GS^* + *µ^GS^*) and Poisson’s ratio *ν^GS^* = *λ^GS^*/2(*λ^GS^* + *µ^GS^*) of the tissue. Additionally, *I*_1_ = tr ***C*** is the first invariant of the right Cauchy-Green tensor ***C*** = ***F****^T^ **F***. The collagen fiber network exhibits viscoelastic behavior, and its time-dependent response is modeled using the reactive viscoelastic framework described in [45]. Following this framework, the collagen fiber network is modeled as a reactive constrained mixture of strong bonds (e.g., covalent bonds) and weak bonds (e.g., hydrogen bonds). Strong and weak bonds are responsible for the equilibrium and instantaneous mechanical responses of the material, respectively. Upon loading, the weak bonds break and reform into a stress-free state, giving rise to the time-dependent mechanical response of the tissue. This bond reaction is governed by mass balance in which the weak bond mass fraction *w* is an observable state variable. The strain energy density of a reactive viscoelastic material is given by:

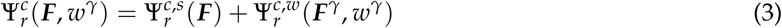

where Ψ*^c^*^,*s*^ and Ψ*^c^*^,*w*^ represent the strong and weak bonds contributions, and ***F****^γ^* and *w^γ^* are the deformation gradient and the mass fraction of the weak bonds. Based on previous work modeling the equilibrium mechanical behavior of cervical tissue [16; 28], strong bonds are modeled as continuous fiber bundles dispersed in 3D about a preferential orientation, 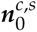, defined in the reference configuration. The strain energy density of the strong bonds is given by [47]:

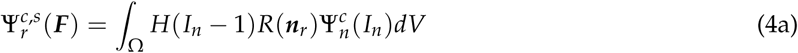

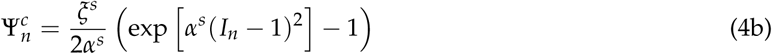

where Ω is a unit sphere over which the integral is calculated, ***n****_r_* = {*n_r_*_1_, *n_r_*_2_, *n_r_*_3_} is the fiber orientation unit vector in the reference configuration that spans all directions from the origin to the unit sphere, and *I_n_* = ***n****_r_* · ***C*** · ***n****_r_* is the strain invariant along the ***n****_r_* direction. *H*(*I_n_* − 1) is a Heaviside step function to include only fibers experiencing tension, and *R*(***n****_r_*) is the fiber density distribution function. The functional 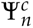 represents the strain energy density of the fiber bundle oriented along the single direction ***n****_r_*. The constants *ξ^s^ >* 0 and *α^s^* ≥ 0 are the initial fiber stiffness and exponential stiffening constant of the strong bonds, respectively. Their effect on the force relaxation curve is shown in Supplementary Figure 1 (a) and (b) in the supplementary material. The collagen fibers in the model are assumed to be dispersed following a three-dimensional von Mises distribution:

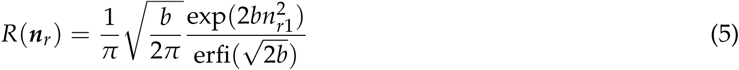

where *I*_0_ is the modified Bessel function of the first kind of order zero, and *b >* 0 is the fiber concentration parameter. Fibers are more dispersed when *b* → 0, and more aligned along the direction 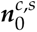 when *b* → ∞. The strain energy density of the weak bonds is given by [47]:

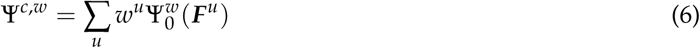

where 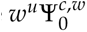 is the strain energy of the weak bond generation *u*, ***F****^u^*(*t*) is the deformation gradient relative of the bond generation *u*, and *w^u^* is the mass fraction satisfying ∑*_u_ w^u^* = 1. The functional 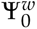 is the strain energy density of a single family of weak bonds when they are all intact, given by:

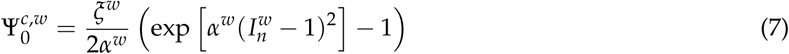

The parameters *ξ^w^ >* 0 and *α^w^* ≥ 0 are the initial fiber stiffness and exponential stiffening constant of the weak bonds, respectively, and 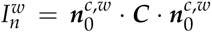, where 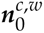 is the weak bonds fiber orientation in the reference configuration. The weak bonds reaction is assumed to be triggered by distortional strains. The influence of the weak bonds’ material parameters on the force relaxation curve is presented in the Supplementary Figure 1 (c) and (d) in the supplementary material. The kinetics of the reaction is controlled by a reduced relaxation function *g*(*t*) that satisfies *g*(0) = 1 and lim*_t_*_→∞_ *g*(*t*) = 0 (see Figure 2 in the supplementary material). We use the continuous relaxation spectrum proposed by Malkin [48]:

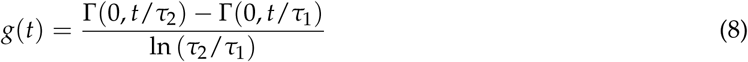

where *τ*_1_ and *τ*_2_ are the characteristic relaxation times and satisfy *τ*_2_ *> τ*_1_ *>* 0, and Γ(*a*, *z*) is the incomplete gamma function. The reduced relaxation function may depend on the strain, such as *g*(*t*, ***F****^u^*(*t*)), enabling the modeling of nonlinear viscoelasticity. Nevertheless, this study found that the strain’s effect on the reduced relaxation function did not substantially enhance our predictions. Consequently, we used a strain-independent version of Eq. 8. The Cauchy stress of the solid mixture is given by:

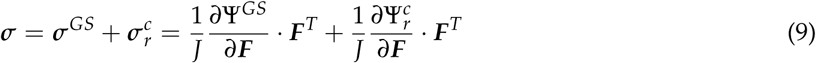

The material behavior of the tissue is determined by eleven (11) material parameters describing the material behavior of the elastic ground matrix (*E^GS^*, *ν^GS^*), the strong bonds of the collagen fiber network (*ξ^s^*, *α^s^* 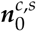, *b*), the weak bonds (*ξ^w^*, *α^w^*, 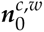), and the reduced relaxation function time constants (*τ*_1_,*τ*_2_). Methods for estimating each of these parameters are described below.

### 2.2 Tissue Collection and Mechanical Testing

To calibrate our constitutive model, we used experimental force relaxation data obtained from uniaxial tension tests, and equilibrium force data from spherical indentation tests on *ex vivo* Rhesus macaque cervical tissue. Details of the tissue collection and mechanical testing protocols can be found in [10]. In summary, the experimental study used cervical tissue specimens from *n_m_* = 12 Rhesus macaques, at four gestational time points: nonpregnant or NP (*n_m_*= 3), early second trimester or E2 (*n_m_* = 3), early third trimester or E3 (*n_m_* = 3), and late third trimester or L3 (*n_m_*= 3), see Figure 1(a). Information from each macaque, including age, gestational age, gravidity, and obstetric history is listed in Table 1. At each gestational stage, sections from the external os (EO) and internal os (IO) of the cervix were cut and split into four quadrants, as shown in Figure 1(b). Quadrant *A* corresponds to the anterior part of the cervix, quadrant *P* is the posterior region, and quadrants *R* and *L* are the cervix’s lateral right and left sides, respectively. From each quadrant, specimens were extracted for mechanical testing.

**Figure 1:**
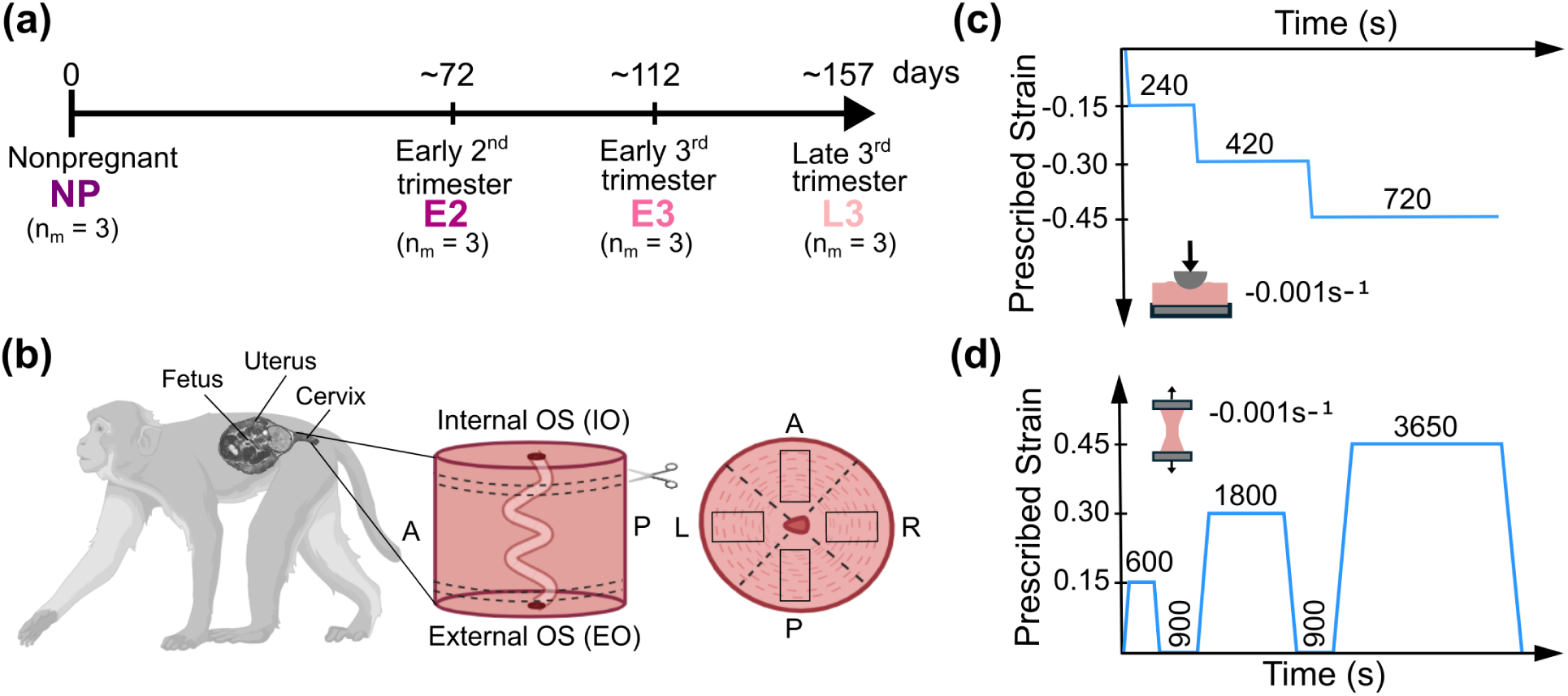
Tissue collection and testing protocols: (a) Cervical tissue samples were collected from *n_m_*= 12 Rhesus macaques at four gestational stages. (b) Specimens were obtained from various anatomical locations, such as the cervix’s external os (EO), internal os (IO), and the anterior (A), posterior (P), right (R), and left (L) sides (adapted from [10]). (c) Uniaxial tension test protocol showing the prescribed strain profile applied to the top grip. (d) Spherical indentation test protocol showing the prescribed strain profile applied to the indenter.

**Table 1:** Summary of cervical specimens collected and used in this study. NP: non-pregnant, E2: early second trimester, E3: early third trimester, L3: Late third trimester, VB: Vaginal birth, CB: cesarean birth, EO: external os, IO: internal os, and N/A: not applicable.

| Macaque ID | Age (yr) | Gestational Group | Gestational Age (days) | Gravidity | Obstetric History | Samples Used | Specimens Analyzed |
| --- | --- | --- | --- | --- | --- | --- | --- |
| NP1 | 15.2 | NP | N/A | 5 | VBx4, CBx1 | 5 | EOx2 IOx3 |
| NP2 | 4.1 | NP | N/A | 0 | N/A | 5 | EOx4, IOx1 |
| NP3 | 11.8 | NP | N/A | 5 | VBx5 | 4 | EOx2, IOx2 |
| E21 | 8.9 | E2 | 71 | 3 | VBx2, CBx1 | 7 | EOx4, IOx3 |
| E22 | 11 | E2 | 73 | 1 | VBx1 | 7 | EOx3, IOx4 |
| E23 | 15.2 | E2 | 72 | 7 | VBx4, CBx3 | 6 | EOx3, IOx3 |
| E31 | 8 | E3 | 110 | 3 | VBx3 | 6 | EOx3, IOx3 |
| E32 | 18 | E3 | 111 | 8 | VBx5, CBx3 | 4 | EOx1, IOx3 |
| E33 | 17.8 | E3 | 116 | 4 | VBx1, CBx3 | 6 | EOx2 IOx4 |
| L31 | 18.4 | L3 | 152 | 2 | VBx1, CBx1 | 6 | EOx3, IOx3 |
| L32 | 9.7 | L3 | 158 | 4 | CBx4 | 7 | EOx3, IOx4 |
| L33 | 19.8 | L3 | 163 | 10 | VBx7, CBx3 | 5 | EOx1, IOx4 |

The mechanical testing protocol for uniaxial tension and indentation testing are shown in Figures 1(c) and (d). Representative force relaxation curves obtained from uniaxial tension and spherical indentation tests are shown in Figures 2(a) - (d). These mechanical tests were conducted using an Instron universal testing machine (Instron, Inc., Norwood, MA) using a 5 N load cell with an accuracy of 0.005 N. The spherical indentation test consisted of a three-level ramp-hold protocol with prescribed indentation depths of 15%, 30%, and 45% of the specimen’s thickness (see Figure 1(c)). We used a spherical indenter of radius *r_ind_* = 1.25 mm. The indenter was held at the same position each cycle for 240, 420, and 720 s to enable tissue relaxation. The force and displacement data over time were recorded.

**Figure 2:**
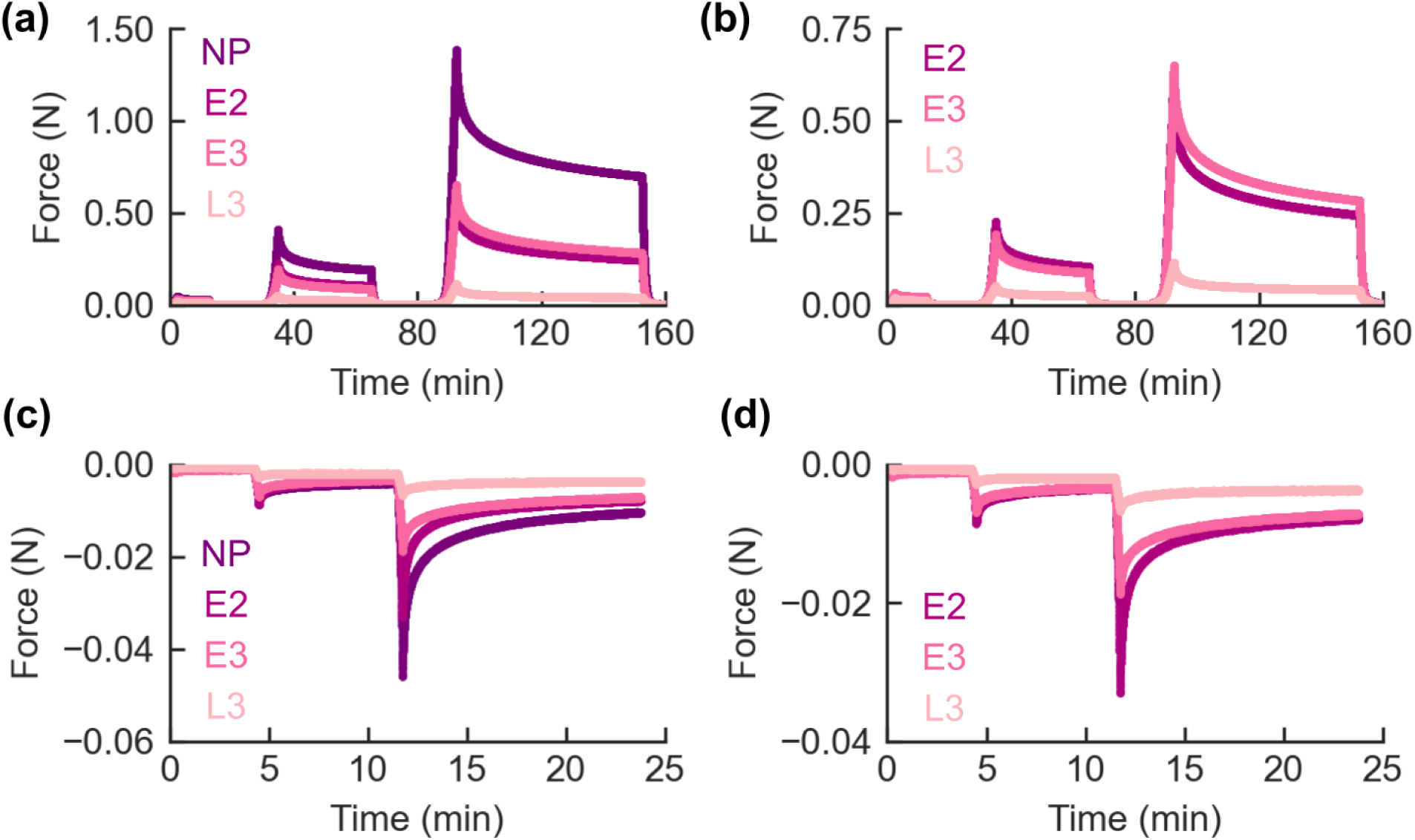
Representative experimental force-relaxation curves obtained from Rhesus macaque cervical specimens at the internal os (IO) tested under (a) - (b) uniaxial tension and (c) - (d) spherical indentation. Curves are presented for cervical tissue at non-pregnancy (NP), early second trimester (E2), early third trimester (E3), and late third trimester (L3).

After indentation, the specimens were mechanically tested under uniaxial tension using a three-level load-hold-unload protocol (see Figure 1(d)). Prescribed strains of 15%, 30%, and 45 % of the grip-to-grip distance were applied. At the hold part of each cycle, the specimen was kept in place for 600, 1800, and 3650 s to allow relaxation. The specimens were left unstressed between cycles for 15 min to allow recovery. Force and displacement data over time were also recorded. For uniaxial tension and indentation, specimens were tested in a fluid solution of phosphate-buffered saline (PBS) and a small amount of 1,2-Ethanedithiol (EDT). The load was applied at an engineering strain rate of 0.1% s^−1^. Following mechanical testing, hysteresis curves for uniaxial tension tests were reviewed. Those that exhibited discontinuities in the loading curve, indicatives of tissue damage or slippage, were excluded from subsequent analysis.

### 2.3 Inverse Finite Element Analysis (IFEA)

To estimate the material parameters of the constitutive model in Section 2.1, an IFEA was conducted using mechanical testing data from uniaxial tension and spherical indentation tests on cervical tissue. For this purpose, FEA models of the tissue specimens were built using the FE software FEB_IO_[49] version 3.7. The geometries and boundary conditions of the models are shown in Figs. 3(a) and (c). The uniaxial tension model consists of a rectangular prism with the dimensions of the real tissue specimens (height *l*_0_, width *w*_0_, and thickness *d*_0_). The geometry is discretized with HEX-8 elements of size *h* = 0.4 mm, chosen based on a sensitivity study to balance accuracy in the force relaxation response and computational efficiency. The model includes a rigid tensile grip to simulate the load application from the tension experiments. An upward displacement *u_z_* is applied according to the experimental protocol shown in Figure 1(a). A tied-elastic contact is used between the grip and the specimen interface. The bottom of the specimen is fixed (*u_x_* = *u_y_* = *u_z_* = 0), and traction-free conditions (***t****_n_* = 0) are applied to the lateral sides of the geometry.

**Figure 3:**
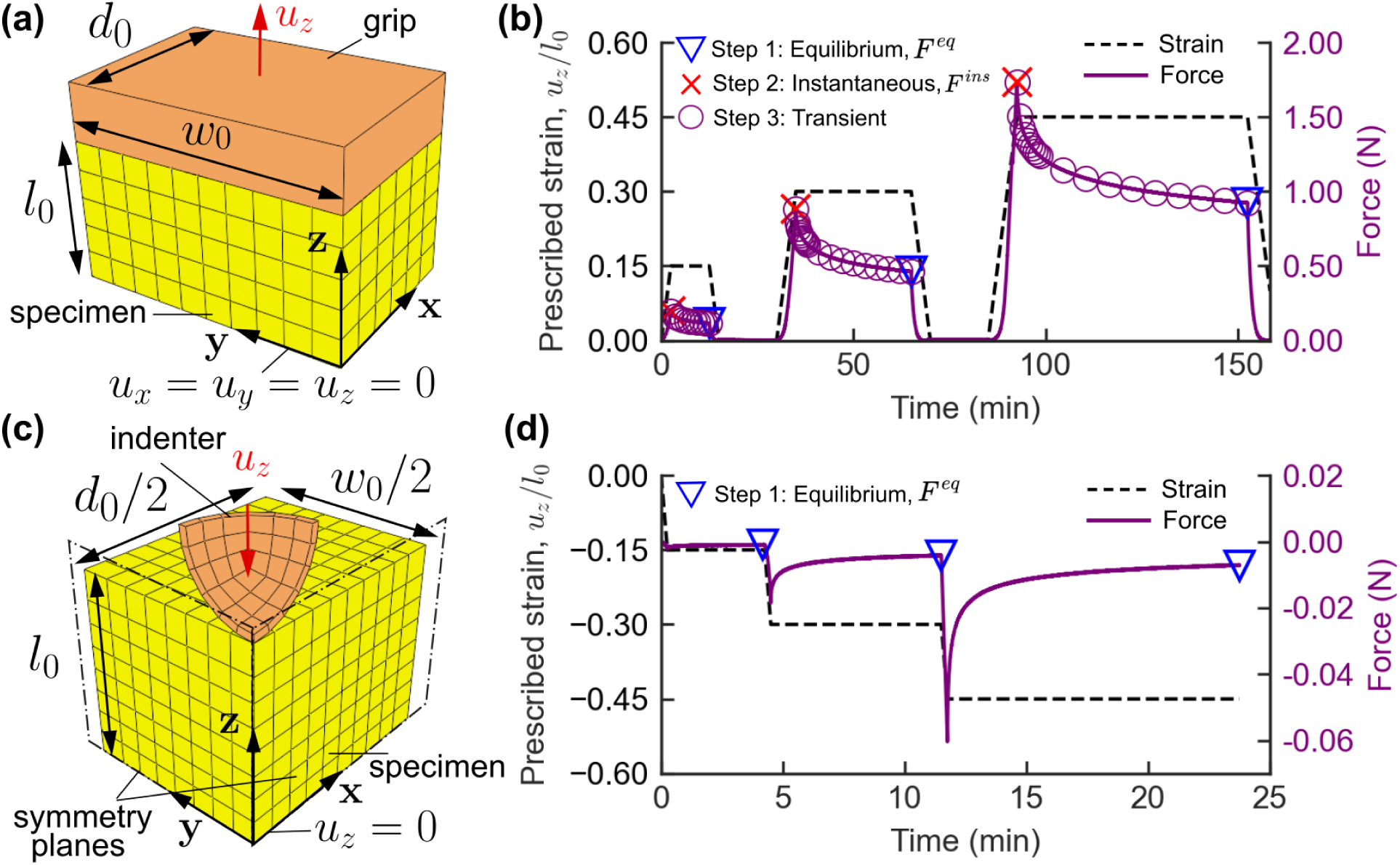
(a) Finite element model of the uniaxial tension test specimen and (b) force-relaxation curve for a non-pregnant cervical specimen extracted from the external os. (c) Finite element quarter model used for simulating the spherical indentation test. (d) Force-relaxation curve from indentation for a non-pregnant cervical specimen from the external os. Blue triangular markers indicate experimental points for IFEA step 1, red cross markers for step 2, and circular markers for step 3.

For simulating the indentation experiment, a quarter FEA model was built with the geometry and boundary conditions shown in Figure 3(b). The geometry consists of a deformable rectangular prism with real specimen thickness *l*_0_, half its length *d*_0_, and half its width *w*_0_. A quarter model of the spherical indenter with radius *r_ind_* = 1.25 mm is included and modeled as a rigid body. Symmetric boundary conditions were applied on two of the lateral sides of the specimen. A frictionless sliding-elastic contact is applied between the top of the specimen and the indenter’s bottom face. The load was applied by prescribing a downward displacement *u_z_* to the indenter based on the experimental loading protocol shown in Figure 1(d). The displacement of the bottom boundary of the specimen was constrained in the direction of the load (*u_z_* = 0). Traction-free conditions (***t****_n_* = 0) are applied at the lateral sides of the specimen, different from the symmetry planes. HEX-8 elements with size *h* = 0.4 mm were used.

To capture the material behavior of the specimens, we used the constitutive model described in Section 2.1. Because collagen fibers in the cervix stroma preferentially align circumferentially around the cervical canal, the preferential fiber direction of the strong bonds, 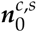, was assumed to be parallel to the **y**-axis for the uniaxial tension model and parallel to the **x**-axis for the indentation model. Weak bonds direction, 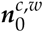, were aligned with the load direction (**z**-axis). The fiber concentration parameter *b* values were taken from a previous study [10]. Eight material parameters (*E^GS^*, *ν^GS^*, *ξ^s^*, *α^s^*, *ξ^w^*, *α^w^*, *τ*_1_, *τ*_2_) were calibrated using the D_AKOTA_ optimization software version 6.17.0 [50]. We used an IFEA approach consisting of three steps. First, we estimated the equilibrium elastic material properties ***θ***_1_ = (*E^GS^*, *ν^GS^*, *ξ^s^*, *α^s^*) by fitting the equilibrium force response, *F^eq^*, of a hyperelastic FEA model to the experimental forces obtained at the last step of each load-hold cycle from indentation and uniaxial tension tests (see Figs. 3(a) and (b)). These forces approximate the equilibrium mechanical response of the tissue. We used a hyperelastic material model consisting of a solid mixture of a neo-hookean ground substance (Eq. 2) and the continuous fiber distribution used for the strong bonds (Eq. 4a). A hybrid sequential optimization approach was employed, starting with the efficient global optimization (EGO) method to identify a promising parameter space region, followed by a nonlinear least-squares method for local refinement.

The second step of the IFEA approach consists of obtaining initial estimates of the weak bonds parameters ***θ***_2_ = (*ξ^w^*, *α^w^*), which capture the instantaneous mechanical behavior of the material. To do so, the hyperelastic model was fit to the experimental instantaneous force response, *F^ins^*, of the three load-hold cycles from uniaxial tension (see Figure 3(a)). In this step, the constitutive model corresponds to a solid mixture of the constitutive model used in step 1 and the exponential fiber constitutive model used for weak bonds (Eq. 7). The same hybrid sequential optimization approach from step 1 was used. Finally, in the third step, the transient force data from the uniaxial tension experiment (see Figure 3(a)) and the initial guesses obtained from steps 1 and 2 were used to calculate the final values for the parameters ***θ***_3_ = (*ξ^s^*, *α^s^*, *ξ^w^*, *α^w^*, *τ*_1_, *τ*_2_). The full anisotropic reactive viscoelastic constitutive model was used in this final step. A nonlinear least-squares method was used to search for the optimum parameter values locally. For all three steps, the material parameters were iteratively optimized with the boundaries listed in Table 2. The quality of each IFEA fit was evaluated using the coefficient of determination (*r*^2^) and the normalized root mean squared error (*NRMSE*) given by:

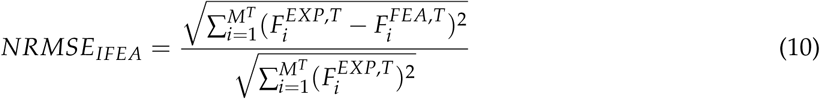

**Table 2:** Boundary values of material parameters used for inverse finite element analysis (IFEA).

| Material Parameter | Symbol | Range |
| --- | --- | --- |
| Young's modulus ground substance | $E^{GS}$ | [0.1 - 100] kPa |
| Poisson's ratio ground substance | $\nu^{GS}$ | [0.0 - 0.499] |
| Initial fiber stiffness strong bonds | $\zeta^s$ | [0.1 - 1000] kPa |
| Stiffening effect constant strong bonds | $\alpha^s$ | $[10^{-3} - 5.0]$ |
| Initial fiber stiffness weak bonds | $\zeta^w$ | [0.1 - 100] kPa |
| Stiffening effect constant weak bonds | $\alpha^w$ | $[10^{-3} - 5.0]$ |
| Reduced relax. function time constant 1 | $\tau_1$ | [0.1 - 100] s |
| Reduced relax. function time constant 2 | $\tau_2$ | [0.1 - 10000] s |

where *M_T_*= 49 is the number of uni-axial tension experimental data points used for parameter optimization, *F^FEA,T^* are the forces obtained from FEA simulations, and *F^EXP,T^* are the experimental forces from uniaxial tension tests.

### 2.4 Material Model Validation

The constitutive material model was validated by comparing the finite element analysis (FEA) simulation results to transient experimental data obtained from the same cervical samples but not used for IFEA.

The FEA predictions and experimental measurements of geometrical changes in the specimens’ width (*w*) and thickness (*d*) in the middle of the sample were compared as a function of time during uniaxial tension testing (Figure 4). For this purpose, the sample width stretch was defined as *λ_w_*(*t*) = *w*(*t*)/*w*_0_ and the thickness stretch *λ_d_* (*t*) = *d*(*t*)/*d*_0_, where *w*_0_ and *d*_0_ were the initial width and thickness of the sample, respectively. To measure the stretch of the specimen cross-section during uniaxial tension tensing, a digital image correlation (DIC) with a custom image segmentation algorithm was used as described in [10]. This algorithm quantified changes in the width and thickness of the uniaxial tension specimens as shown in by creating masks outlining the specimen geometry and averaging the mask in the respective directions (Figures 4 (a) and (c)). Representative experimental curves of the width and thickness stretch over time are shown in Figures 4 (b) and (d). We used the normalized root mean squared error to compute the validation error as follows:

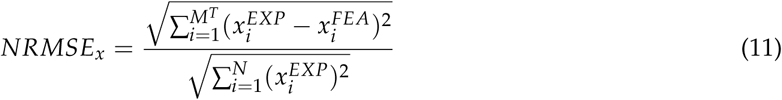

**Figure 4:**
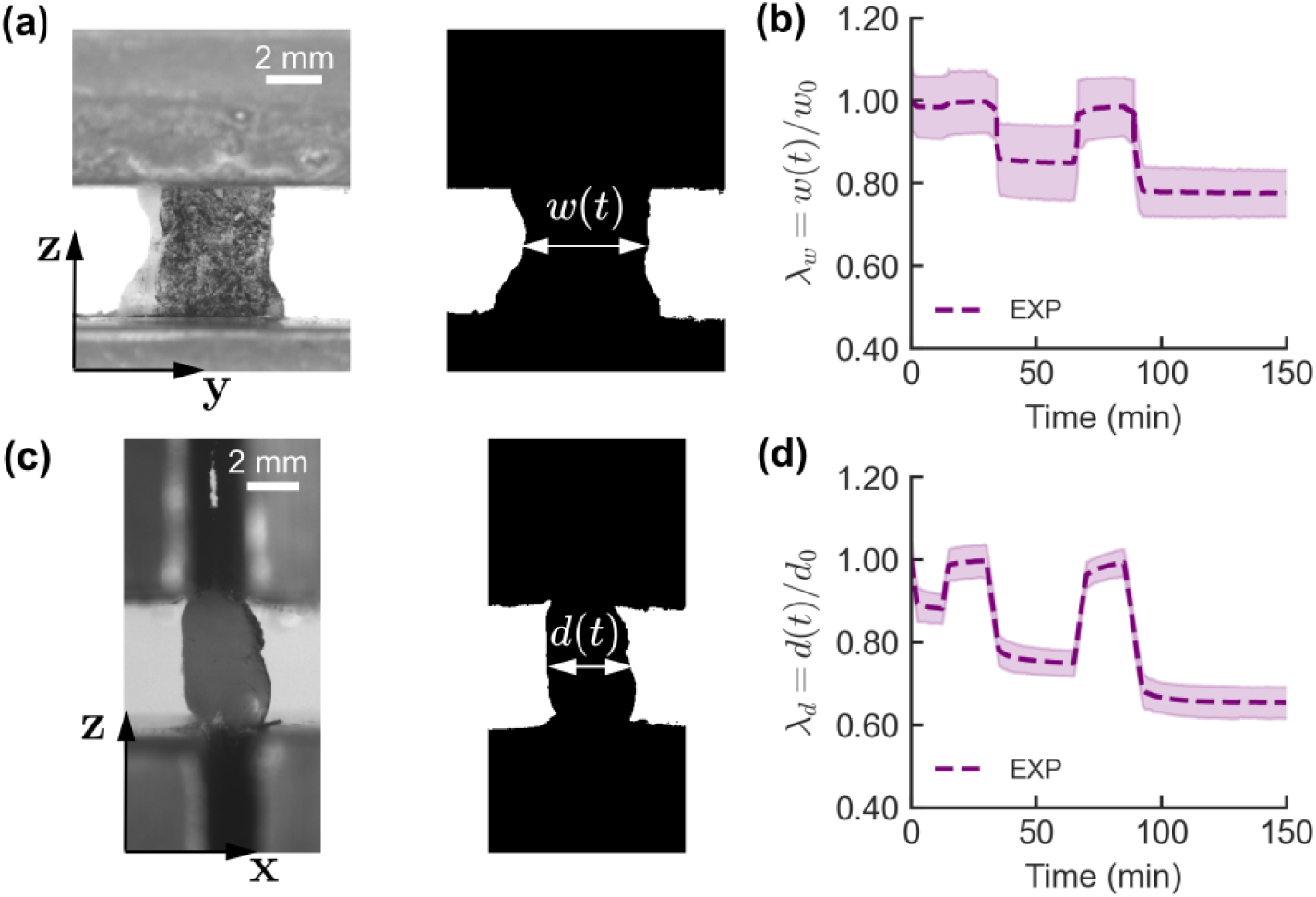
(a) Front view of uniaxial tension cervical specimen obtained from DIC, and mask of the specimen used to obtain the specimen width *w*(*t*) during testing. (b) Width stretch *λ_w_* over time. (c) Lateral view of uniaxial tension cervical specimen obtained from DIC, and mask of the specimen used to obtain the specimen thickness *d*(*t*) as a function of time. (d) Thickness stretch *λ_d_* over time. The shaded regions represent ± 2 times the standard deviation.

where the variable *x* represents either of the stretches *λ_w_* or *λ_d_* obtained from experiments (*EXP*) or finite element simulations (*FEA*).

### 2.5 Statistical Analysis

A linear mixed-effect model assessed statistically significant differences in material parameters across gestational age and anatomical location. The gestational age group and cervical ora (external or internal os) were selected as the model fixed effects based on a sensitivity study. Rhesus macaque ID was set as the random variable to account for material parameters obtained from multiple specimens extracted from the same macaque. A logarithmic transformation was applied to material properties such as Young’s modulus (*E^GS^*), initial fiber stiffness (*ξ^s^*and *ξ^w^*), and relaxation time constants (*τ*_1_, *τ*_2_, and *τ*_1/2_) to satisfy the assumptions of normality as assessed by QQ-plots. No transformations were made for material parameters such as Poisson’s ratio (*ν^GS^*) and the stiffening effect constants (*α^s^* and *α^w^*). Statistical analysis was performed in RS_TUDIO_ version 2023.12 using the _LME_ library. The p-value symbols in the plots follow a standard GraphPad style (ns : *p >* 0.1, ^#^ : *p* ≤ 0.1, ^∗^ : *p* ≤ 0.05, ^∗∗^ : *p* ≤ 0.01, and ^∗∗∗^ : *p* ≤ 0.001).

## 3 Results

### 3.1 Quality of IFEA fits

Representative force relaxation curves comparing results from finite element analysis (FEA) and uniaxial tension experiments (EXP) in specimens at the cervix IO are presented in Figure 5. The FEA simulations were performed using the material parameters obtained from the IFEA procedure described in Section 2.3. As shown in Figure 5, for all gestational groups, the constitutive model formulated in this work captures well the force relaxation response of Rhesus macaque cervical tissue deformed at prescribed strains between 15% and 45%. Table 3 summarizes the mean and standard deviation values of the coefficient of determination (*r*^2^) and IFEA error (*NRMSE_IFEA_*) per gestational group. In general, these results highlight the good agreement between FEA and experimental force-relaxation data (0.03 ≤ NRMSE*_IFEA_* ≤ 0.23 and 0.84 ≤ *r*^2^ ≤ 0.99).

**Figure 5:**
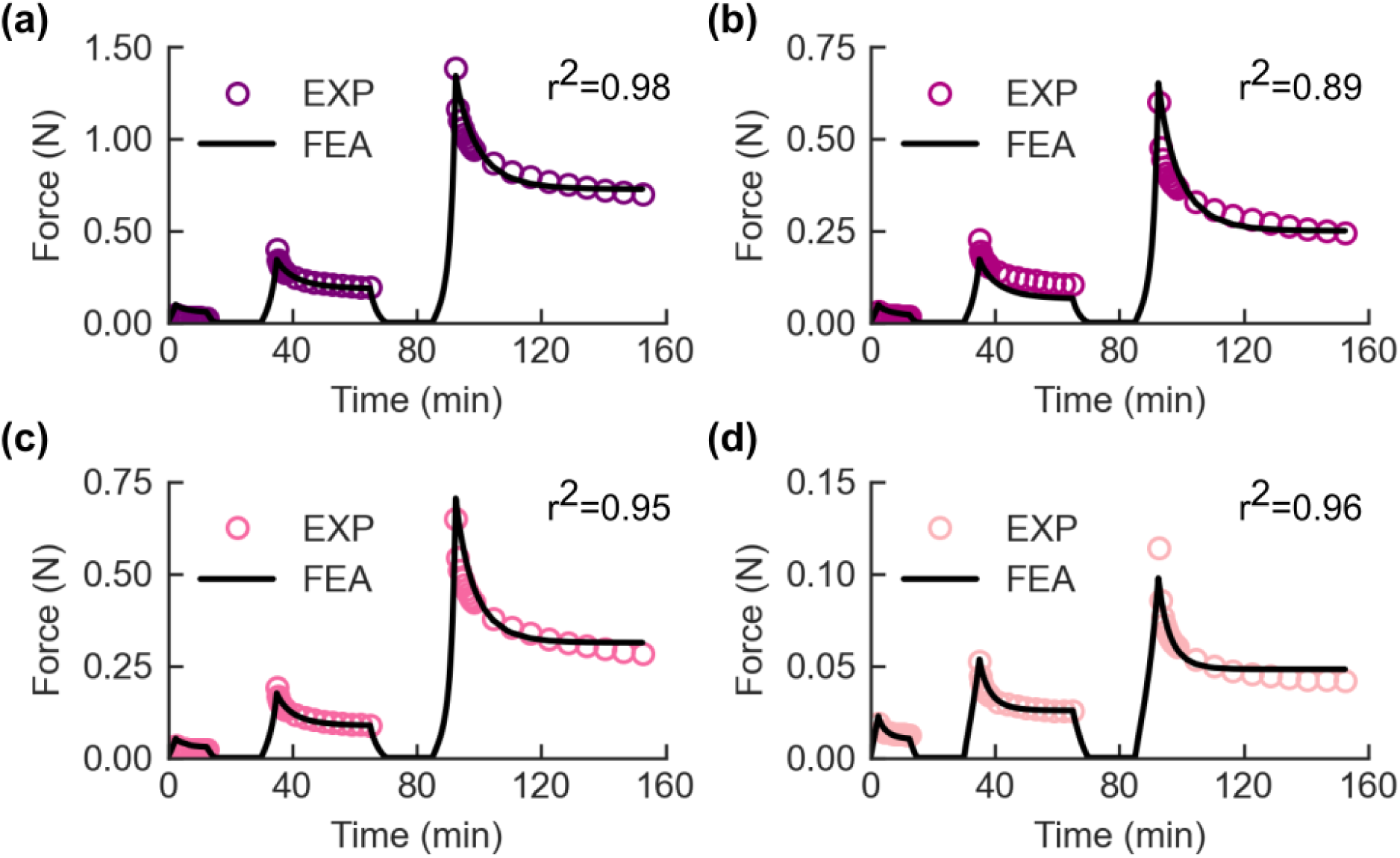
Representative force relaxation curves comparing results from FEA simulations and experimental (EXP) uniaxial tension tests on cervical specimens from Rhesus macaques at different stages of pregnancy: (a) non-pregnant (NP), (b) early second trimester (E2), (c) early third trimester (E3), and (d) late third trimester (L3). All the specimens in this figure were obtained from the internal os (IO) of the cervix.

**Table 3:** Summary of the quality of IFEA fits, highlighting the coefficient of determination (*r*^2^) and the normalized root mean squared error (NRMSE*_IFEA_*). The table shows the average values standard deviations of these quantities for each gestational group under study (NP: non-pregnant, E2: early second trimester, E3: early third trimester, and L3: late third trimester).

| Gestational Group | $r^2$ | $NRMSE_{IFEA}$ |
| --- | --- | --- |
| NP ($n_s = 14$) | $0.970 \pm 0.014$ | $0.105 \pm 0.030$ |
| E2 ($n_s = 20$) | $0.960 \pm 0.024$ | $0.115 \pm 0.041$ |
| E3 ($n_s = 16$) | $0.954 \pm 0.039$ | $0.116 \pm 0.059$ |
| L3 ($n_s = 18$) | $0.967 \pm 0.018$ | $0.088 \pm 0.030$ |

### 3.2 Material Properties of the Macaque Cervix Across Gestation

The average equilibrium engineering stress, 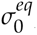, vs. the grip-to-grip prescribed strain *ε_z_* from uniaxial tension tests on Rhesus macaque cervical specimens is presented in Figure 6 (a) for the four gestational groups under study. As can be observed, 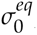 is non-linear, and the equilibrium stiffness of the tissue, measured as the slope of the stress vs. prescribed strain curves, decreases significantly from non-pregnant (NP) to the late third trimester (L3) gestational state. This decrease in stiffness is observed at both low and high strains. The IFEA-fitted material parameters that contribute to the equilibrium stress of Rhesus macaque cervical tissue for each gestational group are shown in Figures 6 (b)–(e). The average and standard deviation values of these parameters are listed in Table 4 for each gestational group. In terms of the material parameters of the ground substance (GS), the average values of Young’s modulus (*E^GS^*) varied from 1.63 to 4.57 kPa, with no statistically significant changes across gestation (Figure 6(c)). The mean values of Poisson’s ratio (*ν^GS^*) were found to range between 0.16 and 0.20, with no notable statistically significant differences across gestational stages (Figure 6(d)).

**Figure 6:**
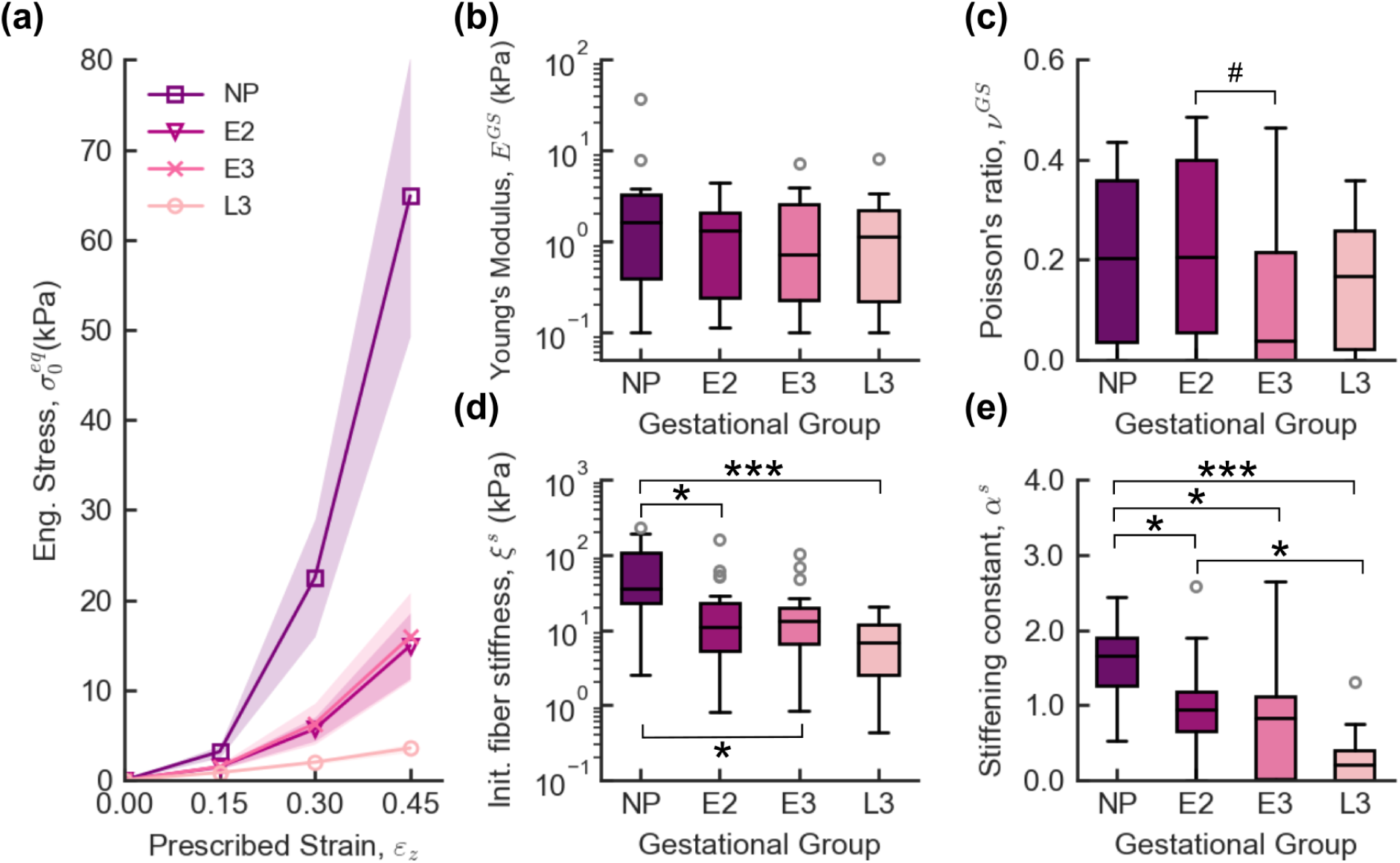
(a) Mean s.m.e of the nominal equilibrium stress 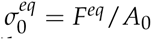 vs prescribed grip-to-grip strain *ε_z_*, and variations across gestation in material properties of Rhesus macaque cervical tissue from (b)-(c) the ground substance and (d)-(e) the strong bonds. Gestational groups, NP: non-pregnancy, E2: early second trimester, E3: early third trimester, and L3: late third trimester. Statistical significance: ^#^*<* 0.1; * *<* 0.05; ** *<* 0.01; *** *<* 0.005.

**Table 4:** Best-fit material parameters obtained from IFEA for Rhesus macaque cervical specimens at the different gestation groups under study. Material parameters include Young’s modulus (*E^GS^*) and Poisson’s ratio (*ν^GS^*) of the ground substance, initial fiber stiffness (*ξ^s^*) and stiffening constant (*α^s^*) of the strong bonds, initial fiber stiffness (*ξ^w^*) and stiffening constant *α^w^* of the weak bonds, and the reduced relaxation function time constants (*τ*_1_,*τ*_2_). Format: average ± standard deviation.

| Gestational Age | $E^{GS}$ (kPa) | $\nu^{GS}$ | $\xi^s$ (kPa) | $\alpha^s$ |
| --- | --- | --- | --- | --- |
| NP ($n_s = 14$) | $4.57 \pm 9.58$ | $0.20 \pm 0.16$ | $73.05 \pm 71.66$ | $1.56 \pm 0.55$ |
| E2 ($n_s = 20$) | $1.59 \pm 1.43$ | $0.23 \pm 0.18$ | $24.51 \pm 36.88$ | $0.97 \pm 0.62$ |
| E3 ($n_s = 16$) | $1.60 \pm 2.03$ | $0.14 \pm 0.17$ | $22.56 \pm 28.34$ | $0.73 \pm 0.74$ |
| L3 ($n_s = 18$) | $1.63 \pm 1.97$ | $0.16 \pm 0.13$ | $7.59 \pm 5.57$ | $0.27 \pm 0.34$ |
| Gestational Age | $\xi^w$ (kPa) | $\alpha^w$ | $\tau_1$ (s) | $\tau_2$ (s) |
| NP ($n_s = 14$) | $18.11 \pm 20.21$ | $1.56 \pm 0.84$ | $27 \pm 28$ | $700 \pm 366$ |
| E2 ($n_s = 20$) | $3.98 \pm 5.32$ | $2.05 \pm 0.76$ | $40 \pm 33$ | $468 \pm 139$ |
| E3 ($n_s = 16$) | $5.17 \pm 9.18$ | $1.66 \pm 1.19$ | $49 \pm 42$ | $537 \pm 240$ |
| L3 ($n_s = 18$) | $1.24 \pm 1.20$ | $1.73 \pm 0.89$ | $32 \pm 29$ | $438 \pm 170$ |

IFEA results show significant variations in the initial collagen fiber stiffness (*ξ^s^*) and stiffening constant (*α^s^*) of the strong bonds of the Rhesus macaque cervix during pregnancy (Figures 6(d)–(e)). These two material properties characterize the equilibrium tensile stiffness of the tissue under low and high strains, respectively. As shown in Figure 6(b), the median values of *ξ^s^* decrease with advancing gestational age. Specifically, L3 specimens exhibit an initial stiffness approximately ten times lower than the NP specimens, with a statistically significant difference (p = 0.004). Additionally, there are marginally significant reductions in *ξ^s^* from NP to E2 (p = 0.04) and from NP to E3 (p = 0.03). No significant differences in *ξ^s^* are detected between the E2, E3, and L3 groups. These changes in *ξ^s^*can be better observed in Figure 6 (a), where notable differences in the mean equilibrium stress 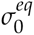 at the low strain region (less than 30% strain) are observed between the NP and L3 groups.

The stiffening constant *α^s^* also decreases with gestational status. The median of *α^s^* is significantly lower for the L3 gestational group than for the NP group (p = 0.001). Some significant differences in *α^s^* were also observed between the pairs NP - E2 (p = 0.04), NP - E3 (p = 0.01), and E2 - L3 (p = 0.02). These differences in *α^s^*suggest that the equilibrium stiffness of the Rhesus macaque cervical tissue at high levels of deformation decreases significantly during pregnancy. Overall, the trends observed in the material properties *ξ^s^* and *α^s^* indicate that under tensile deformation and quasi-static loading conditions, the Rhesus macaque cervix becomes significantly more compliant and extensible as pregnancy progresses.

Experimental curves of the average instantaneous engineering stress, 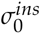, vs. prescribed grip-to-grip strain, *ε_z_*, for the different gestational groups under study are shown in Figure 7(a). Furthermore, Figures 7(c) - (d) show the results of the material parameters that contribute to the instantaneous and time-dependent response of the Rhesus macaque cervical tissue as a function of gestational status. The means and standard deviations of these material parameters for each gestational group are shown in Table 4. Material parameters associated with the collagen weak bonds are shown in Figures 7(a) and (b). A significant decrease in the initial fiber stiffness, *ξ^w^*, of the weak bonds is observed from the NP to the L3 gestational status (p = 0.001). Significant differences in *ξ^w^* are also observed between NP and E2 groups (p = 0.011) as well as E2 to L3 groups (p = 0.013), suggesting a considerable decrease in the instantaneous stiffness of the Rhesus macaque cervix during pregnancy. This result can also be observed in Figure 7(a), where notable differences in the instantaneous stress-strain response are shown between NP and L3 groups. No statistically significant differences are observed in the material parameter *α^w^*.

**Figure 7:**
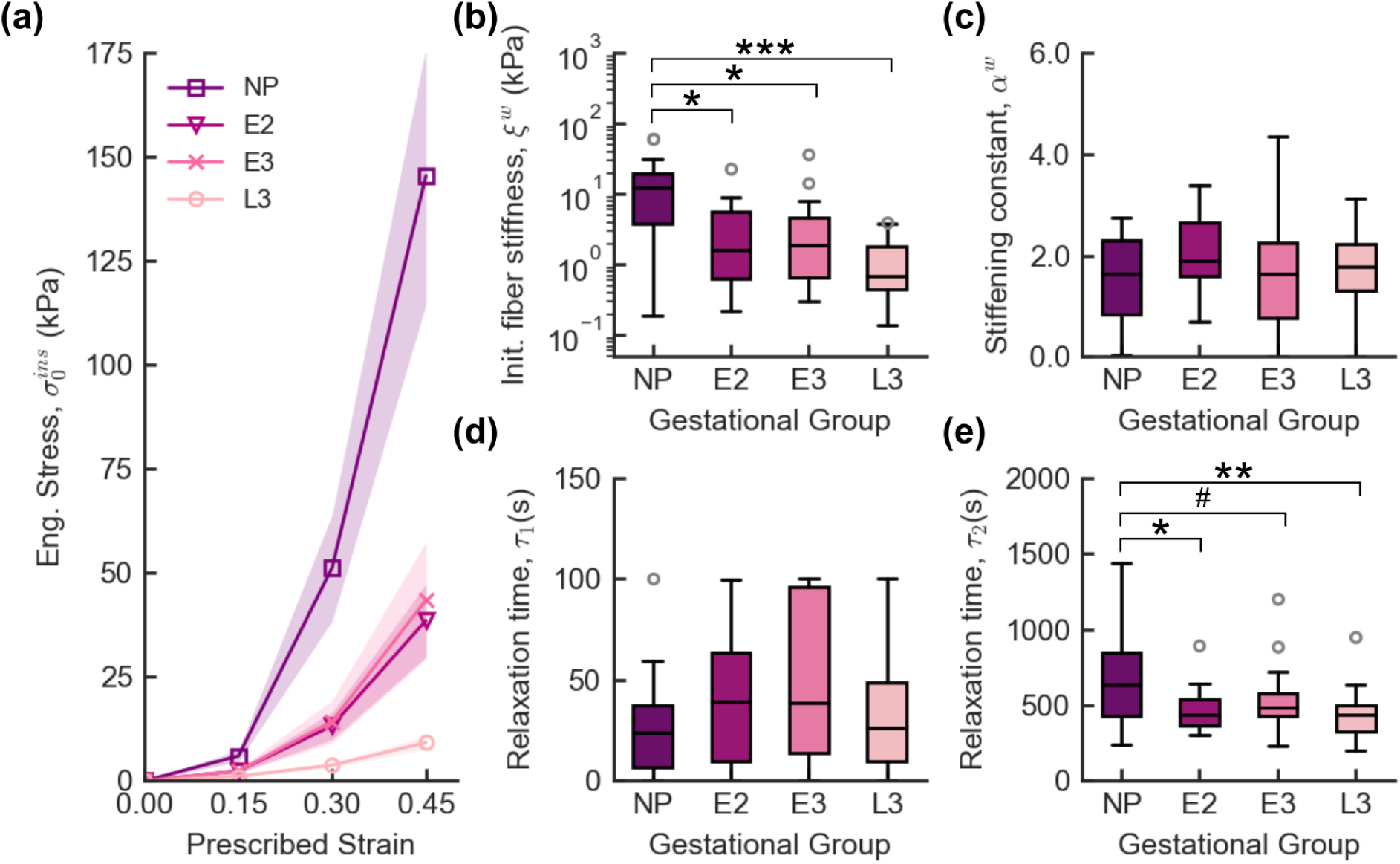
(a) Mean s.m.e of the engineering instantaneous stress, 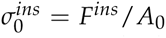, vs. prescribed grip-to-grip strain *ε_z_*, and (b) - (e) variations in the weak bonds material properties and relaxation constants across gestation. Gestational groups, NP: non-pregnancy, E2: early second trimester, E3: early third trimester, and L3: late third trimester. Statistical significance: ^#^*<* 0.1; * *<* 0.05; ** *<* 0.01; *** *<* 0.005.

The results of the relaxation time constants *τ*_1_ and *τ*_2_ used in the continuous reduced relaxation function in Eq. 8 are shown in Figures 7(c) and (d). Significant differences were observed in the relaxation time *τ*_2_ between NP and L3 (p = 0.009), NP and E3 (p = 0.07), and NP and E2 (p=0.02) gestational groups, indicating alterations in the relaxation behavior of Rhesus macaque cervical tissue from non-pregnancy to late pregnancy. To better understand the differences in the relaxation behavior of Rhesus macaque cervical tissue across gestation, we calculated the relaxation half-time (*τ*_1/2_) for each load-hold-unload cycle in the uniaxial tension tests. *τ*_1/2_ denotes the time during the hold period when the force decreases by half the difference between its peak or instantaneous (*F^ins^*) and equilibrium (*F^eq^*) or end of the hold force values. The greater *τ*_1/2_ is, the more time it takes for the tissue to reach equilibrium. This quantity is calculated for each load-hold-unload cycle with prescribed strains of 15%, 30%, and 45%. The results of the relaxation half-time are shown in Figure 8. At 45% strain, the relaxation time in L3 specimens was significantly lower compared to E3 (p = 0.06) and NP specimens (p = 0.09), see Figure 8(a). Moreover, for the NP, E2, and E3 specimens, *τ*_1/2_ increases significantly (p *<* 0.001) with the level of prescribed strain, indicating that macaque cervical tissue takes more time to relax to equilibrium as the level of prescribed tensile strain increases (see Figures 8(b), (c), and (d)). Interestingly, for the L3 group, *τ*_1/2_ increases from 0.15 to 0.30 but remains constant from 0.3 to 0.45, as shown in Figure 8(e). These results suggest that for prescribed strains greater than 0.3, the relaxation of late-pregnant tissue becomes strain-independent.

**Figure 8:**
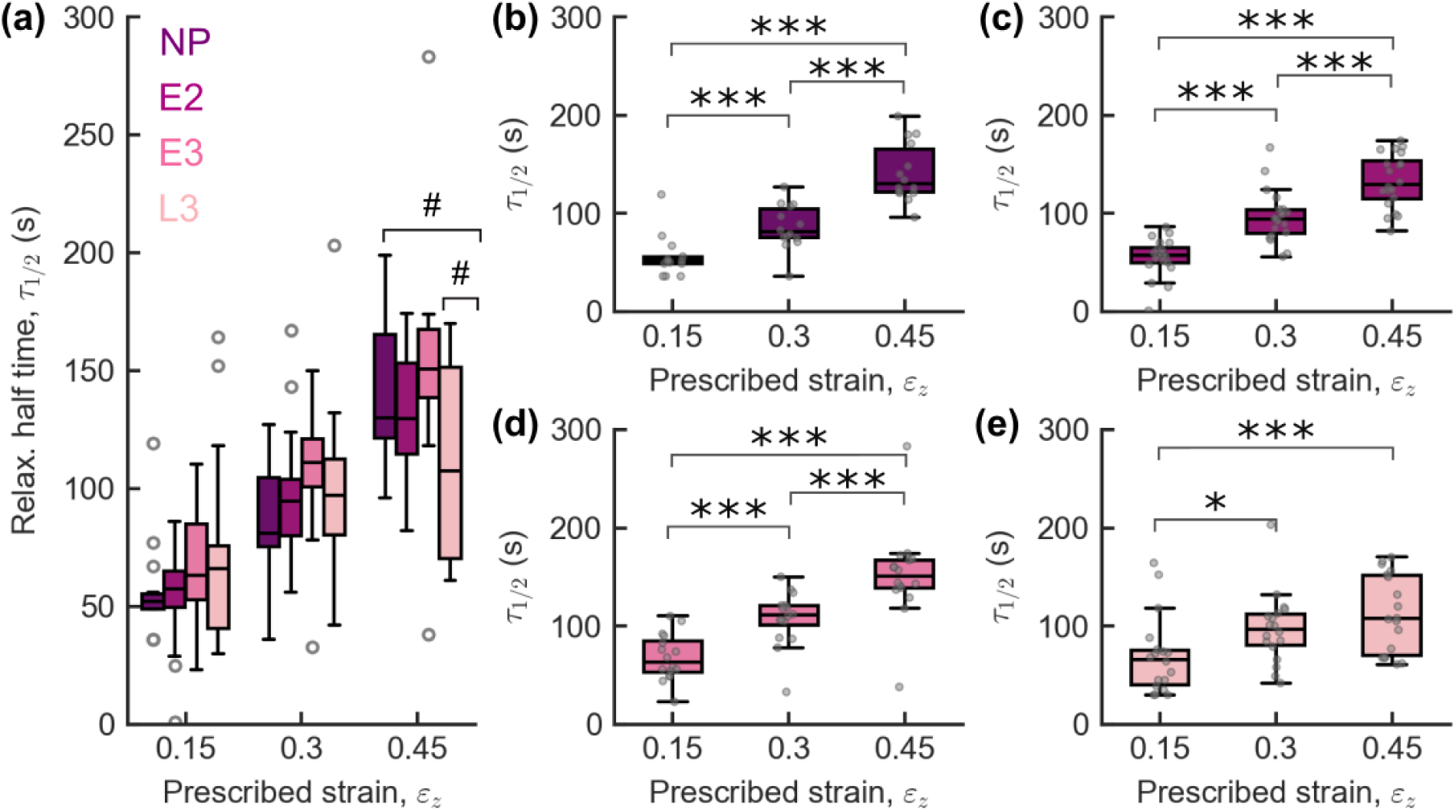
(a) Results relaxation half-time *τ*_1/2_ as a function of prescribed grip-to-grip strain *ε_z_*for all cervical specimens. Relaxation halt-time *τ*_1/2_ as a function prescribed strain for cervical specimens obtained at (b) non-pregnancy (NP), (c) early second trimester (E2), (d) early third trimester (E3), and (d) late third trimester (L3). Statistical significance: ^#^*<* 0.1; * *<* 0.05; ** *<* 0.01; *** *<* 0.005.

### 3.3 Material Properties at Different Anatomical Locations

This section analyzes the differences in material properties of the macaque cervix at various anatomical locations, including the internal os (IO), external os (EO), and different quadrant regions (anterior, posterior, left, and right). Figure 9 compares the initial fiber stiffness *ξ^s^* and stiffening constant *α^s^*of the strong bonds obtained from the specimens at the internal os (IO) and external os (EO) for the different gestational groups under study. Values of fiber stiffness *ξ^s^* are significantly higher at IO compared to the EO, particularly for the E2 (p=0.003) and L3 (p = 0.005) gestational groups. A similar trend is observed for the material parameter *α^s^* in the second trimester (p = 0.07), but with a lower level of statistical significance. These results suggest that the equilibrium stiffness of Rhesus macaque cervical tissue decreases at a higher rate at the EO compared to the IO. No significant differences were observed in other material properties between EO and IO and as a function of the quadrant region. A summary of the results of material properties as a function of ora is shown in Supplementary Table 1.

**Figure 9:**
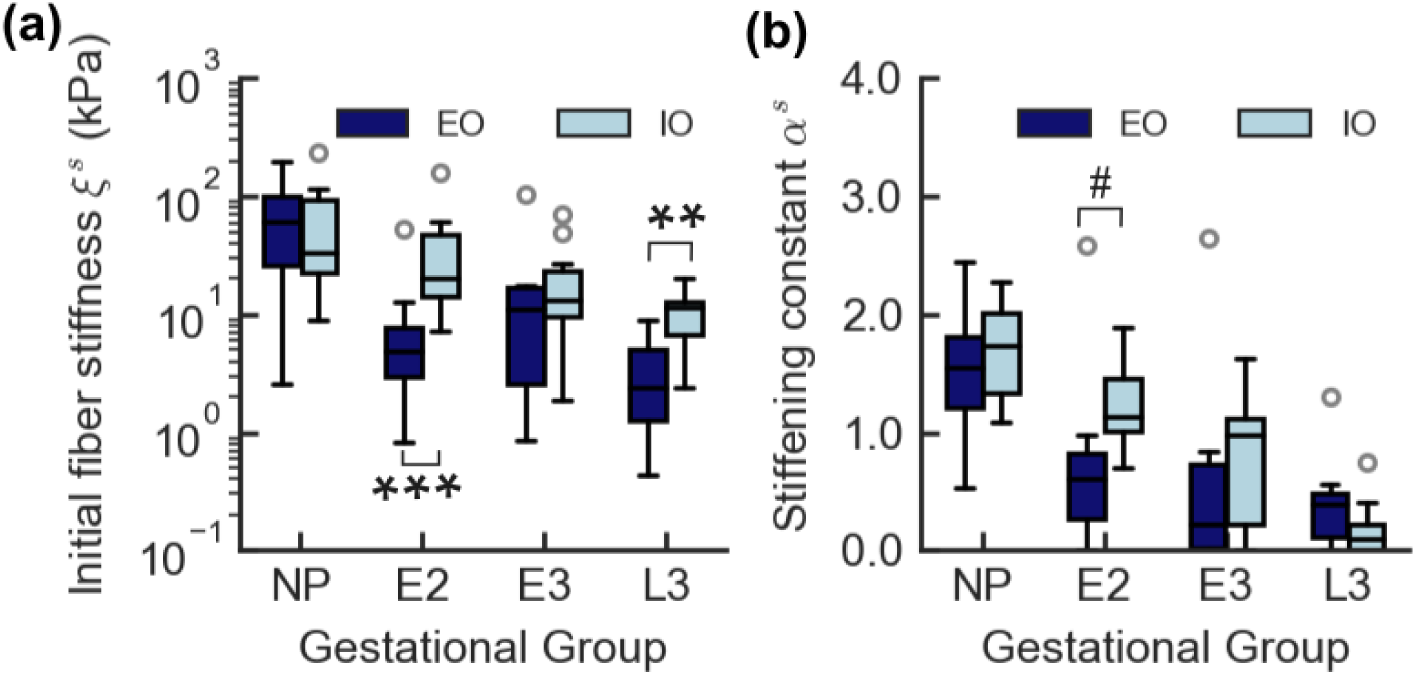
Results reactive viscoelastic material parameters obtained at the cervix external os (EO) and internal os (IO) as a function of gestational status. NP: non-pregnancy, E2: early second trimester, E3: early third trimester, and L3: late third trimester. Statistical significance: ^#^*<* 0.1; * *<* 0.05; ** *<* 0.01; *** *<* 0.005.

### 3.4 Material Model Validation

A summary of the material model validation results for each gestational group is presented in Table 5. Representative curves of the transient width and thickness stretch ratios, *λ_w_*and *λ_d_*, obtained from the segmented masks and FEA simulations, are shown in Figure 10. On average, the model captures well the behavior of cervical tissue under tension (NRMSE*_λw_ <* 0.083 and NRMSE*_λd_ <* 0.142). The constitutive model better captures the transient response of the width stretch than the thickness stretch. In some cases, the constitutive model underestimates the transient depth at prescribed strains above 30% (see Figure 10(b)). We believe these discrepancies arise from tissue swelling and time-dependent tissue behavior influenced by poroelasticity, which the current constitutive model does not account for. In uniaxial tension experiments, the specimen’s transverse dimensions contract due to the Poisson effect. Under this compressive state of stress in the transversal direction, the time-dependent behavior of the tissue could be significantly affected by poroelasticity.

**Figure 10:**
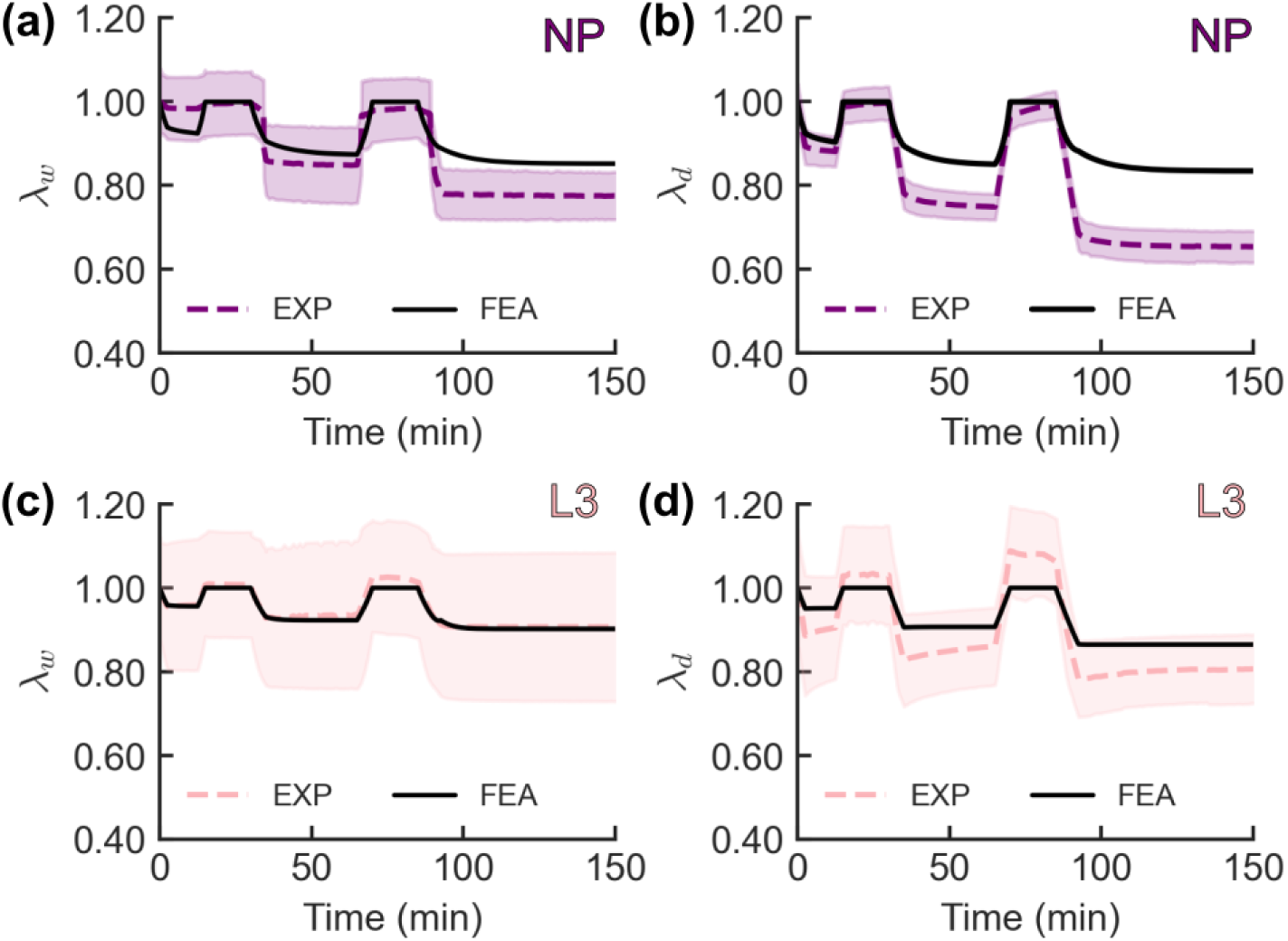
Representative curves of (a) width stretch *λ_w_* and (b) thickness stretch *λ_d_* over time for a non-pregnant (NP) specimen. Representative curves of (c) width stretch *λ_w_* and (d) thickness stretch *λ_d_* over time for a later third trimester (L3) cervical specimen. Curves were obtained from experiments (EXP) and finite element simulation (FEA). The shaded regions represent ± 2 times the standard deviation.

**Table 5:** Summary of material model validation results of the specimens’ width *λ_w_* and thickness *λ_d_* stretch. Format: average ± standard deviation.

| Gestational Age | NRMSE $\lambda_w$ | NRMSE $\lambda_d$ |
| --- | --- | --- |
| NP ($n_s = 14$) | $0.053 \pm 0.031$ | $0.116 \pm 0.091$ |
| E2 ($n_s = 20$) | $0.067 \pm 0.051$ | $0.106 \pm 0.087$ |
| E3 ($n_s = 16$) | $0.083 \pm 0.056$ | $0.142 \pm 0.106$ |
| L3 ($n_s = 18$) | $0.076 \pm 0.059$ | $0.125 \pm 0.089$ |

## 4 Discussion

In this study, the biomechanics of cervical remodeling of the Rhesus macaque was characterized. The biomechanics was characterized by quantifying changes in the tensile time-dependent material properties of *ex vivo* cervical tissue obtained at four relevant gestational points, from non-pregnancy to the late third trimester and at various anatomical locations. For this purpose, an anisotropic reactive viscoelastic constitutive model inspired by the solid components of the cervical ECM was formulated and calibrated. Compared to other viscoelastic models for large deformations, the reactive viscoelastic framework is advantageous because it models the complex anisotropic viscoelastic behavior of cervical tissue as a molecular process. This process involves the breakdown and reformation of weak non-covalent bonds within the collagen fiber network, consistent with the molecular interpretation of intrinsic viscoelasticity [51; 24]. The material model parameters were calibrated using an IFEA approach, which minimizes the differences between force relaxation data from uniaxial tension experiments and FEA simulations. In general, the constitutive model formulated in this study fits the uniaxial tension relaxation response well at prescribed strain levels ranging from 15 % to 45 % (*r*^2^ = 0.962 ± 0.026 and *NRMSE_IFEA_*= 0.106 ± 0.042). Furthermore, it captures well the transient changes in the specimens’ crossectional area (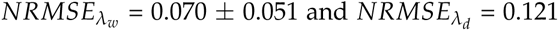 ± 0.092)

Across gestation, the Rhesus macaque cervical tissue’s equilibrium and time-dependent material properties showed drastic changes. In particular, a statistically significant decrease in the initial fiber stiffness *ξ^s^* and stiffening constant *α^s^*of the strong bonds from non-pregnancy (NP) to the late third trimester (L3) was observed. These two material properties affect the nonlinear equilibrium stiffness of the tissue at low and high strains, respectively. The decrease in these two material properties suggests that the cervix of Rhesus macaques becomes more extensible and compliant under tensile deformation as pregnancy progresses. This equilibrium cervical tissue compliance change is observed in the stress-strain curves in Figure 6(a). These results in equilibrium fiber stiffness throughout gestation are consistent with trends observed in previous studies in *ex vivo* cervical tissue from Rhesus macaques, mice, and humans [10; 17; 28; 12]. We hypothesize that the increase in compliance and extensibility with gestational age is attributed to both an increase in the disorganization of the collagen fiber network and a decrease in the quantity of intermolecular mature collagen crosslinks, as observed in the murine cervix [14; 30; 52].

There is also a significant decrease in the weak bonds initial fiber stiffness, *ξ^w^*, throughout pregnancy. This material parameter affects the instantaneous elastic modulus of the tissue. The stiffening effect constant of the weak bonds *α^w^* remained unchanged throughout gestation. These results suggest that the instantaneous cervical stiffness of the Rhesus macaque cervical tissue decreases considerably from the non-pregnant status to the late third trimester. The decrease in the instantaneous initial fiber stiffness is particularly relevant for shear wave elastastrography imaging (SWEI) applications that subject cervical tissue to low-strain and high-strain rate deformation, which does not allow the tissue to relax to equilibrium. SWEI studies in macaque cervical tissue have shown that shear wave speed (SWS), a measure of instantaneous tissue stiffness, decreases significantly during pregnancy[18; 44]. No significant changes in the equilibrium material properties of the ground substance were observed with gestation, which agrees with trends observed in previous quasi-static studies in Rhesus macaques [10]. However, equilibrium material properties of the ground substance did change throughout gestation in tensile studies of mouse cervical tissue [28]. These results could be attributed to differences in remodeling patterns between the two species and experimental protocols.

The intrinsic viscoelasticity of the Rhesus macaque cervix was also found to be affected by gestational status. The relaxation time constant *τ*_2_ significantly decreased throughout gestation, particularly from the non-pregnant to the late third-trimester status. By analyzing the half-relaxation time *τ*_1/2_, a measure of how viscous the tissue is, at each load-hold-unload uniaxial tension cycles, it was found that the relaxation time is strain-dependent for the NP, E2, and E3 groups and significantly increases with the level of applied strain. Interestingly, for L3 tissue, the relaxation half time *τ*_1/2_ is strain-independent at prescribed grip-to-grip strains between 30 and 45% and lower than for the other gestational ages at 45% prescribed strain. Similar trends were observed in a previous study using cervical tissue from rats [36]. In this study, late third trimester cervical tissue relaxes faster than non-pregnant, second trimester, and early third trimester tissue. We believe this fast recovery to equilibrium and lower stiffness of the late third trimester cervix allows it to stretch considerably and dissipate tensile hoop stresses that will be induced by the fetus during labor faster and more efficiently, preventing its damage and/or rupture. These results highlight the potential of using the stress relaxation time constant as a biomarker of cervical remodeling. Previous studies have shown that the stress relaxation time of late pregnant murine cervical tissue increases due to the lack of proteoglycans (PGs) such as decorin and biglycan [16]. Additionally, relaxation time has been found to increase in the cervices of patients with cervical insufficiency (CI) compared to healthy patients [33]. Therefore, the viscoelastic relaxation time may provide insights into abnormal cervical remodeling, potentially helping predict the onset of sPTB.

There are material property differences between the internal (IO) and external os (EO), but not between the posterior and anterior sides of the cervix. The strong bonds median initial fiber stiffness *ξ^s^*and stiffening constant *α^s^* from tissue dissected from the IO were significantly higher than those of the EO, mainly in the early second (E2) and late third trimesters (L3). Additionally, the rate of decrease in *ξ^s^* and *α^s^* with gestation was higher for the EO specimens than for the IO specimens. No significant differences were observed in other material properties with respect to the anatomical location. Rosado et al. [44] found significant differences in SWS measurements from SWEI in *ex vivo* unripe cervical specimens from Rhesus macaques obtained at the EO and IO. They observed no significant differences in SWS based on anatomical location in ripened cervical tissue [44]. Furthermore, SWS measurements in both *in vivo* and *ex vivo* tissues were larger on the posterior side of the cervix than on its anterior side [18; 44]. Histological analysis revealed that the human and murine cervix exhibit a 3D gradient of smooth muscle cells (SMC) from the IO to the EO[7; 53]. The human pregnant cervix contains up to 60% SMCs at the IO compared to 20% at the EO [7]. The gravid mouse cervix comprises up to 80% SMCs at the IO, compared to 20% SMCs at the EO [53]. Because the cervical specimens tested in this study were subjected to freeze-thaw cycles, SMC destruction disabled active material properties, which may be present *in vivo*. Nevertheless, the limited and uneven number of specimens obtained from different regions of the cervix makes it difficult to assess any anatomical changes in cervical stiffness and viscoelasticity of rhesus macaques in this study.

There are some limitations related to the modeling approach, sample size, and the IFEA method in this study. Previous studies have highlighted the asymmetry in the mechanical response of the cervix from compression and tension owing to the activation of different deformation mechanisms from the collagen fiber network, ground substance, and their interaction with interstitial fluid [9; 12; 13]. The constitutive material model presented here captures the uniaxial force relaxation response of rhesus macaque cervical tissue well. Under uniaxial tension, deformation mechanisms associated with the collagen fiber network, such as fiber uncrimping, alignment, sliding, and breakdown of weak crosslinks, contribute to the time-dependent mechanical response of the cervix [23; 24; 25]. While the material model effectively accounts for the cervical quasi-equilibrium mechanical response from spherical indentation, it does not capture cervical tissue compressive time-dependent mechanical behavior. During uniaxial deformation, the cross-sectional area of the specimens is subjected to compressive stresses due to Poisson’s effect, so deformation mechanisms such as poroelasticity become relevant. Furthermore, time-dependent mechanical behavior may arise from the ground substance, which is not accounted for in the model. PGs and GAGs can indirectly affect the tensile viscoelastic response of tissues by interacting with collagen fibers and swelling the collagen network, thereby increasing fiber dispersion[54]. Although GAGs such as HA are known to increase significantly in cervical tissue near the onset of labor[32; 55], it has yet to be determined how HA affects the time-dependent mechanical response of cervical tissue under tensile loading. In our future work, we will focus on understanding the effect of interactions between the ground substance and the collagen fiber network in the viscoelasticity of the cervix.

There are also limitations related to the specimen sample size that are inherently associated with studies on non-human primates such as the Rhesus macaque. The current study analyzed cervical specimens from three monkeys per gestational period. A minimum of five cervical specimens per macaque were used to obtain material parameters. As a result, it was particularly difficult to assess differences in material properties with anatomical location, for example, among the different sides of the cervix: anterior, posterior, and left/right sides. Nevertheless, the sample size was good enough to observe significant differences in the material properties across gestation. Lastly, as the IFEA procedure involves optimizing eight distinct material parameters, it cannot guarantee the uniqueness of our solution. However, our hybrid optimization approach, which involves both global and local optimization methods, performs better than traditional optimization algorithms.

## 5 Summary and conclusions

In this work, we quantitatively study the cervical remodeling of the Rhesus macaque cervix through changes in its *ex vivo* tensile material properties across gestation. An anisotropic reactive viscoelastic constitutive model of cervical tissue is formulated, inspired by the solid components of the cervical ECM. The model simulates the time-dependent response of cervical tissue based on the theory of reactive constrained mixtures, in which viscoelasticity results from a molecular reaction involving the breakdown and reformation of weak non-covalent bonds (e.g., hydrogen bonds) within the collagen fiber network. The material model parameters were calibrated with experimental force relaxation data from uniaxial tension experiments and equilibrium force data from spherical indentation tests obtained in cervical specimens at four distinct and relevant gestational points. The Rhesus macaque cervical tissue’s equilibrium and material properties showed drastic changes throughout gestation. We observed a statistically significant decrease in the initial fiber stiffness of the strong and weak bonds, responsible for the equilibrium and instantaneous stiffness of the tissue, and increased cervical extensibility from the non-pregnant to the late third trimester status. Furthermore, we observed a significantly lower relaxation time in cervical tissue from the late third trimester compared to the other gestational groups, particularly at prescribed tensile strains of 45%. Interestingly, the stress relaxation time of late third trimester tissue remained constant for prescribed strains between 30 and 45%, while it significantly increased with strain for the other gestational groups. These results highlight how the intrinsic viscoelasticity of cervical tissue is correlated to structural changes in the cervical ECM involved in its remodeling during pregnancy and its potential to characterize normal/abnormal cervical remodeling.

## Data availability

Experimental data will be made available on Columbia University’s Academic Commons. A link will be provided upon journal acceptance. Files used for FE simulations are available upon request.

## Acknowledgements

This research was supported by the Eunice Kennedy Shriver National Institute Of Child Health & Human Development under Award Number R01HD091153 to KMM and R01HD072077 to KMM, HF, and TJH. The content is solely the authors’ responsibility and does not necessarily represent the official views of the National Institutes of Health.

## Supplementary Material

**Supplementary Figure 1:**
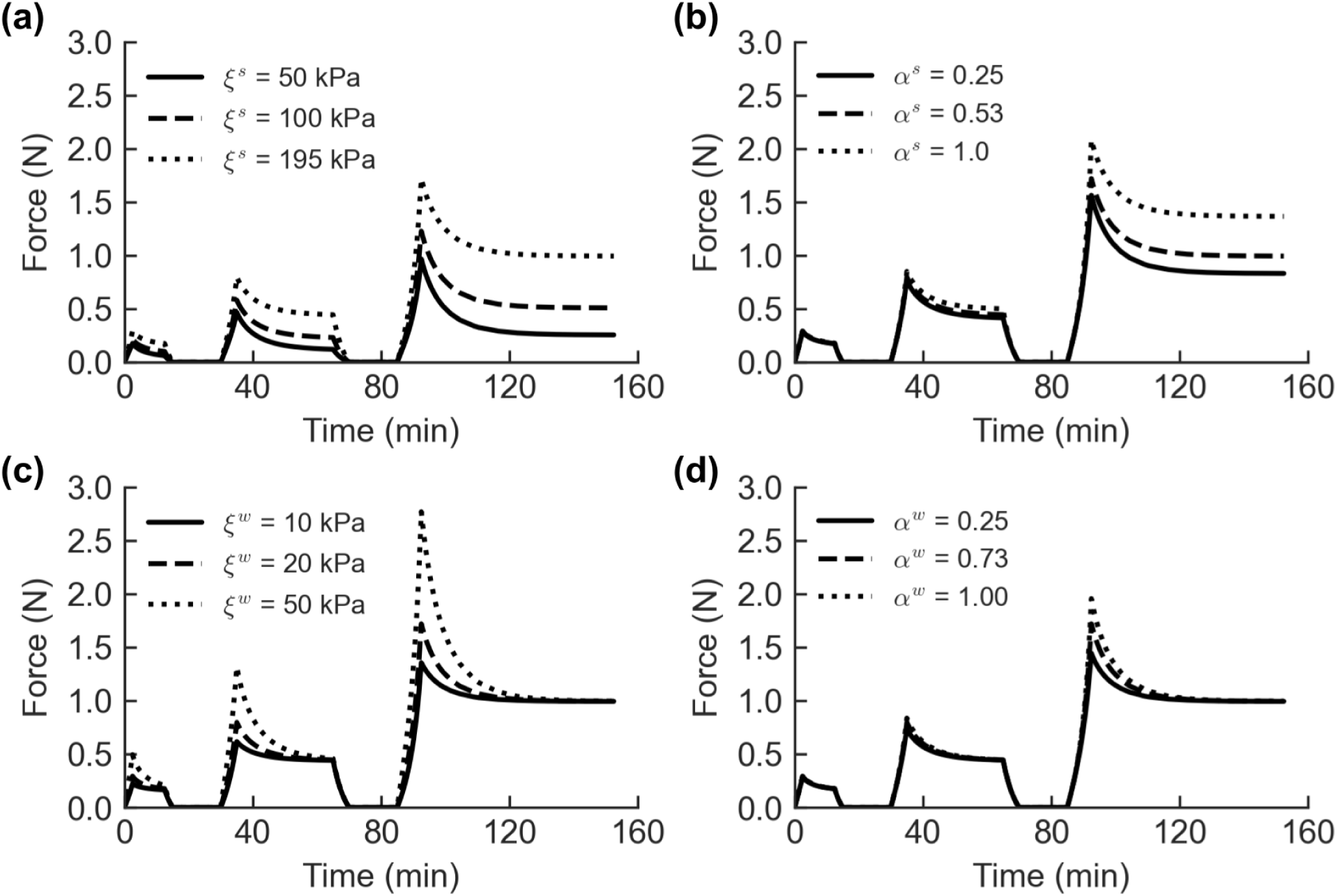
Effect of material parameters on the tensile force relaxation response for a non-pregnant cervical specimen taken from the external os. Strong bonds (a) initial fiber stiffness (*ξ^s^*) and (b) stiffening constant (*α^s^*). Weak bonds (c) initial fiber stiffness (*ξ^w^*) and (d) stiffening constant (*α^w^*). In each subfigure one material parameter is varied while keeping the others fixed, with reference values set at *E^GS^* = 0.187 kPa, *ν^GS^* = 0.09, *ξ^s^* = 195 kPa, *α^s^* = 0.53, *ξ^w^* = 20 kPa, *α^w^* = 0.73, *τ*_1_ = 6.1, *τ*_2_ = 682, and *b* = 1.49.

**Supplementary Figure 2:**
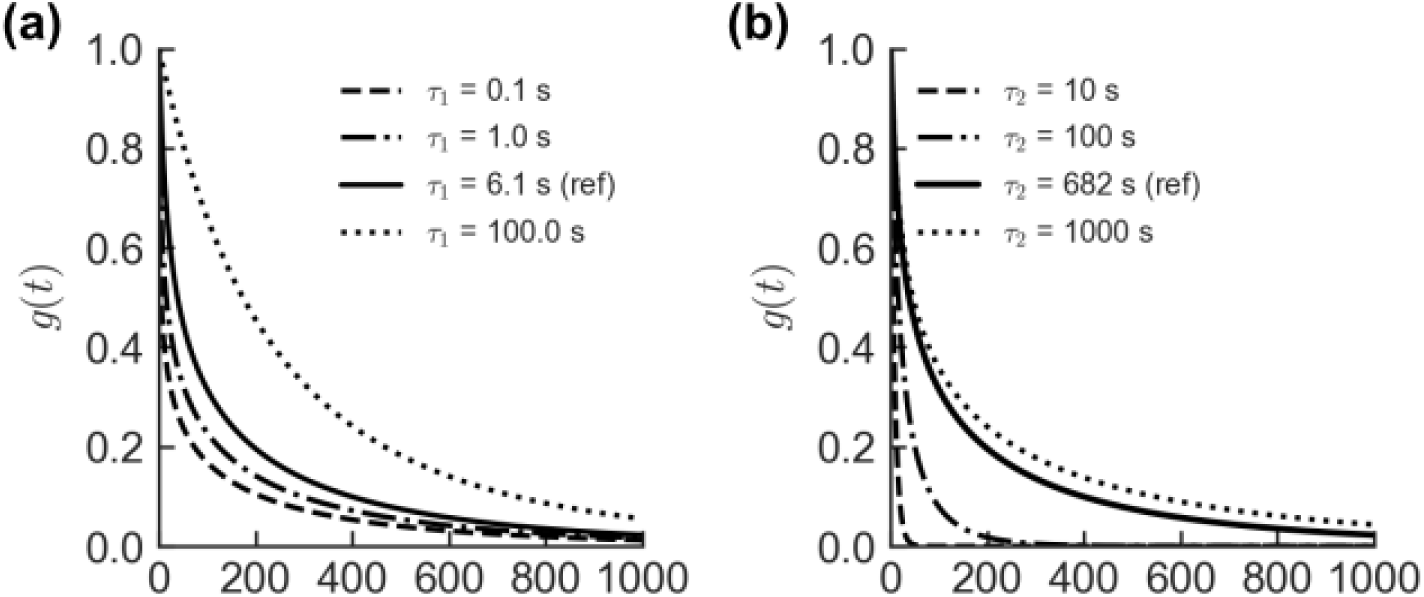
Effect of relaxation constants (a) *τ*_1_ and (b) *τ*_2_ on the reduced relaxation function *g*(*t*). In each subfigure one material parameter is varied while keeping the others fixed, with reference values set at *τ*_1_ = 6.1 and *τ*_2_ = 682.

**Supplementary Table 1:**
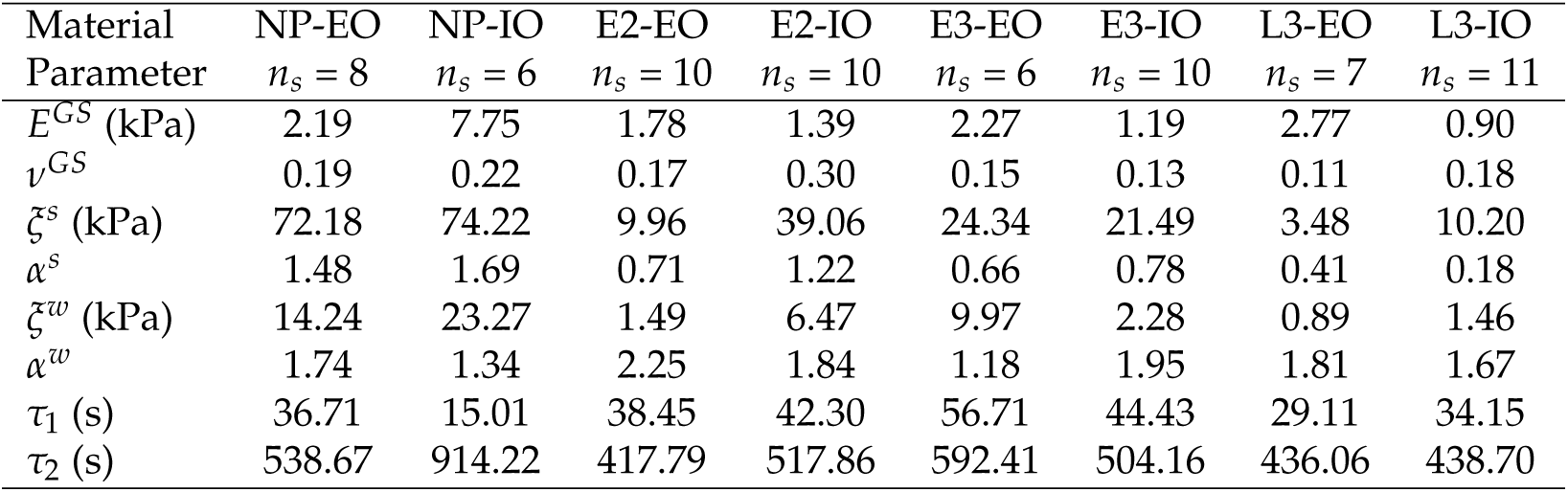
Mean material parameters obtained from IFEA for Rhesus macaque cervical specimens at the different gestation groups under study and anatomical locations, external os (EO) and interna os (IO). Material parameters include Young’s modulus (*E^GS^*) and Poisson’s ratio (*ν^GS^*) of the ground substance, initial fiber stiffness (*ξ^s^*) and stiffening constant *α^s^* of the strong bonds, initial fiber stiffness (*ξ^w^*) and stiffening constant *α^w^* of the weak bonds, and the reduced relaxation function time constants (*τ*_1_,*τ*_2_).

## References

[1] K. M. Myers, H. Feltovich, E. Mazza, J. Vink, M. Bajka, R. J. Wapner, T. J. Hall, and M. House, “The mechanical role of the cervix in pregnancy,” Journal of Biomechanics, vol. 48, no. 9, pp. 1511–1523, 2015. Reproductive Biomechanics.

[2] F. Martyn, F. McAuliffe, and M. Wingfield, “The role of the cervix in fertility: is it time for a reappraisal?,” Human Reproduction, vol. 29, pp. 2092–2098, 07 2014.

[3] A. Xholli, G. Simoncini, S. Vujosevic, G. Trombetta, A. Chiodini, M. F. Ferraro, and A. Cagnacci, “Menstrual pain and elasticity of uterine cervix,” Journal of Clinical Medicine, vol. 10, no. 5, 2021.

[4] B. Timmons, M. Akins, and M. Mahendroo, “Cervical remodeling during pregnancy and parturition,” Trends in Endocrinology & Metabolism, vol. 21, pp. 353–361, June 2010.

[5] C. P. Read, R. A. Word, M. A. Ruscheinsky, B. C. Timmons, and M. S. Mahendroo, “Cervical remodeling during pregnancy and parturition: molecular characterization of the softening phase in mice,” Reproduction, vol. 134, no. 2, pp. 327 – 340, 2007.

[6] M. House, D. L. Kaplan, and S. Socrate, “Relationships between mechanical properties and extracellular matrix constituents of the cervical stroma during pregnancy,” Seminars in Perinatology, vol. 33, no. 5, pp. 300–307, 2009. The Cervix: Prediction and Prevention of Preterm Delivery.

[7] J. Y. Vink, S. Qin, C. O. Brock, N. M. Zork, H. M. Feltovich, X. Chen, P. Urie, K. M. Myers, T. J. Hall, R. Wapner, J. K. Kitajewski, C. J. Shawber, and G. Gallos, “A new paradigm for the role of smooth muscle cells in the human cervix,” American Journal of Obstetrics and Gynecology, vol. 215, pp. 478.e1–478.e11, Oct. 2016.

[8] J. Vink, V. Yu, S. Dahal, J. Lohner, C. Stern-Asher, M. Mourad, G. Davis, Z. Xue, S. Wang, K. Myers, J. Kitajewski, X. Chen, R. J. Wapner, C. V. Ananth, M. Sheetz, and G. Gallos, “Extracellular Matrix Rigidity Modulates Human Cervical Smooth Muscle Contractility—New Insights into Premature Cervical Failure and Spontaneous Preterm Birth,” Reproductive Sciences, vol. 28, pp. 237–251, Jan. 2021.

[9] K. Myers, A. Paskaleva, M. House, and S. Socrate, “Mechanical and biochemical properties of human cervical tissue,” Acta Biomaterialia, vol. 4, no. 1, pp. 104–116, 2008.

[10] S. Fang, L. Shi, J.-S. Y. Vink, H. Feltovich, T. J. Hall, and K. M. Myers, “Equilibrium Mechanical Properties of the Nonhuman Primate Cervix,” Journal of Biomechanical Engineering, vol. 146, p. 081001, 03 2024.

[11] K. Myers, S. Socrate, D. Tzeranis, and M. House, “Changes in the biochemical constituents and morphologic appearance of the human cervical stroma during pregnancy,” European Journal of Obstetrics & Gynecology and Reproductive Biology, vol. 144, pp. S82–S89, May 2009.

[12] K. M. Myers, S. Socrate, A. Paskaleva, and M. House, “A Study of the Anisotropy and Tension/Compression Behavior of Human Cervical Tissue,” Journal of Biomechanical Engineering, vol. 132, p. 021003, 01 2010.

[13] L. Shi and K. Myers, “A finite porous-viscoelastic model capturing mechanical behavior of human cervix under multi-step spherical indentation,” Journal of the Mechanical Behavior of Biomedical Materials, vol. 143, p. 105875, 2023.

[14] K. Yoshida, H. Jiang, M. Kim, J. Vink, S. Cremers, D. Paik, R. Wapner, M. Mahendroo, and K. Myers, “Quantitative evaluation of collagen crosslinks and corresponding tensile mechanical properties in mouse cervical tissue during normal pregnancy,” PLOS ONE, vol. 9, pp. 1–11, 11 2014.

[15] K. Yoshida, C. Jayyosi, N. Lee, M. Mahendroo, and K. M. Myers, “Mechanics of cervical remodelling: insights from rodent models of pregnancy,” Interface Focus, vol. 9, p. 20190026, Oct. 2019.

[16] N. Lee, L. Shi, M. Colon Caraballo, S. Nallasamy, M. Mahendroo, R. V. Iozzo, and K. Myers, “Mechanical Response of Mouse Cervices Lacking Decorin and Biglycan During Pregnancy,” Journal of Biomechanical Engineering, vol. 144, p. 061009, June 2022.

[17] L. Shi, L. Hu, N. Lee, S. Fang, and K. Myers, “Three-dimensional anisotropic hyperelastic constitutive model describing the mechanical response of human and mouse cervix,” Acta Biomaterialia, vol. 150, pp. 277–294, Sept. 2022.

[18] I. M. Rosado-Mendez, L. C. Carlson, K. M. Woo, A. P. Santoso, Q. W. Guerrero, M. L. Palmeri, H. Feltovich, and T. J. Hall, “Quantitative assessment of cervical softening during pregnancy in the rhesus macaque with shear wave elasticity imaging,” Physics in Medicine & Biology, vol. 63, p. 085016, apr 2018.

[19] C. Jayyosi, N. Lee, S. Madhukaran, S. Nallasamy, M. Mahendroo, and K. Myers, “The swelling behavior of the mouse cervix: Changing kinetics with osmolarity and the role of hyaluronan in pregnancy,” Acta Biomaterialia, vol. 135, pp. 414–424, Nov. 2021.

[20] J. Vink and H. Feltovich, “Cervical etiology of spontaneous preterm birth,” Seminars in Fetal and Neonatal Medicine, vol. 21, no. 2, pp. 106–112, 2016. Prediction and prevention of preterm birth and its sequelae.

[21] J.-K. Suh and M. R. DiSilvestro, “Biphasic Poroviscoelastic Behavior of Hydrated Biological Soft Tissue,” Journal of Applied Mechanics, vol. 66, pp. 528–535, June 1999.

[22] M. L. R. Harkness and R. D. Harkness, “Changes in the physical properties of the uterine cervix of the rat during pregnancy,” The Journal of Physiology, vol. 148, no. 3, pp. 524–547, 1959.

[23] B. J. Rigby, N. Hirai, J. D. Spikes, and H. Eyring, “The mechanical properties of rat tail tendon,” Journal of General Physiology, vol. 43, pp. 265–283, 11 1959.

[24] S. Nam, K. H. Hu, M. J. Butte, and O. Chaudhuri, “Strain-enhanced stress relaxation impacts nonlinear elasticity in collagen gels,” Proceedings of the National Academy of Sciences, vol. 113, no. 20, pp. 5492–5497, 2016.

[25] O. Chaudhuri, J. Cooper-White, P. A. Janmey, D. J. Mooney, and V. B. Shenoy, “Effects of extracellular matrix viscoelasticity on cellular behaviour,” Nature, vol. 584, pp. 535–546, Aug. 2020.

[26] A. F. Mak, “Unconfined compression of hydrated viscoelastic tissues: A biphasic poroviscoelastic analysis,” Biorheology, vol. 23, no. 4, pp. 371–383, 1986. PMID: 3779062.

[27] A. J. Grodzinsky and E. H. Frank, Fields, forces, and flows in biological systems. London: Garland science, 2011.

[28] K. Yoshida, M. Mahendroo, J. Vink, R. Wapner, and K. Myers, “Material properties of mouse cervical tissue in normal gestation,” Acta Biomaterialia, vol. 36, pp. 195–209, 2016.

[29] S. Badir, E. Mazza, R. Zimmermann, and M. Bajka, “Cervical softening occurs early in pregnancy: characterization of cervical stiffness in 100 healthy women using the aspiration technique,” Prenatal Diagnosis, vol. 33, no. 8, pp. 737–741, 2013.

[30] M. L. Akins, K. Luby-Phelps, R. A. Bank, and M. Mahendroo, “Cervical Softening During Pregnancy: Regulated Changes in Collagen Cross-Linking and Composition of Matricellular Proteins in the Mouse1,” Biology of Reproduction, vol. 84, pp. 1053–1062, 05 2011.

[31] J. C. Ramella-Roman, M. Mahendroo, C. Raoux, G. Latour, and M.-C. Schanne-Klein, “Quantitative Assessment of Collagen Remodeling during a Murine Pregnancy,” ACS Photonics, vol. 11, pp. 3536–3544, Sept. 2024. Publisher: American Chemical Society.

[32] Y. Akgul, R. Holt, M. Mummert, A. Word, and M. Mahendroo, “Dynamic Changes in Cervical Glycosaminoglycan Composition during Normal Pregnancy and Preterm Birth,” Endocrinology, vol. 153, pp. 3493–3503, 07 2012.

[33] W. A. van Duyl, A. T. M. van der Zon, and A. C. Drogendijk, “Stress relaxation of the human cervix: a new tool for diagnosis of cervical incompetence,” Clinical Physics and Physiological Measurement, vol. 5, p. 207, aug 1984.

[34] W. R. Barone, A. J. Feola, P. A. Moalli, and S. D. Abramowitch, “THE EFFECT OF PREGNANCY AND POSTPARTUM RECOVERY ON THE VISCOELASTIC BEHAVIOR OF THE RAT CERVIX,” Journal of Mechanics in Medicine and Biology, vol. 12, p. 1250009, Mar. 2012.

[35] W. Yao, K. Yoshida, M. Fernandez, J. Vink, R. J. Wapner, C. V. Ananth, M. L. Oyen, and K. M. Myers, “Measuring the compressive viscoelastic mechanical properties of human cervical tissue using indentation,” Journal of the Mechanical Behavior of Biomedical Materials, vol. 34, pp. 18–26, June 2014.

[36] A. Ashofteh Yazdi, A. Esteki, and M. M. Dehghan, “Determination of an average quasi-linear viscoelastic model for the mechanical behavior of rat cervix,” *Proceedings of the Institution of Mechanical Engineers*, Part L: Journal of Materials: Design and Applications, vol. 233, pp. 924–929, May 2019.

[37] R. J. DeWall, T. Varghese, M. A. Kliewer, J. M. Harter, and E. M. Hartenbach, “Compression-dependent viscoelastic behavior of human cervix tissue,” Ultrasonic Imaging, vol. 32, no. 4, pp. 214–228, 2010. PMID: 21213567.

[38] L. Peralta, G. Rus, N. Bochud, and F. Molina, “Assessing viscoelasticity of shear wave propagation in cervical tissue by multiscale computational simulation,” Journal of Biomechanics, vol. 48, pp. 1549–1556, June 2015.

[39] A. Callejas, J. Melchor, I. H. Faris, and G. Rus, “Viscoelastic model characterization of human cervical tissue by torsional waves,” Journal of the Mechanical Behavior of Biomedical Materials, vol. 115, p. 104261, 2021.

[40] M. Mahendroo, “Cervical remodeling in term and preterm birth: insights from an animal model,” REPRODUCTION, vol. 143, no. 4, pp. 429 – 438, 2012.

[41] C. Casteleyn and J. Bakker, “Anatomy of the rhesus monkey (¡em¿macaca mulatta¡/em¿): The essentials for the biomedical researcher,” in Updates on Veterinary Anatomy and Physiology (C. S. Rutland and S. A. El-Gendy, eds.), ch. 1, Rijeka: IntechOpen, 2021.

[42] E. S. E. Hafez and S. Jaszczak, “Comparative anatomy and histology of the cervix uteri in non-human primates,” Primates, vol. 13, pp. 297–314, Sept. 1972.

[43] M. G. Gravett, S. S. Witkin, G. J. Haluska, J. L. Edwards, M. J. Cook, and M. J. Novy, “An experimental model for intraamniotic infection and preterm labor in rhesus monkeys,” American Journal of Obstetrics and Gynecology, vol. 171, no. 6, pp. 1660–1667, 1994.

[44] I. M. Rosado-Mendez, M. L. Palmeri, L. C. Drehfal, Q. W. Guerrero, H. Simmons, H. Feltovich, and T. J. Hall, “Assessment of structural heterogeneity and viscosity in the cervix using shear wave elasticity imaging: Initial results from a rhesus macaque model,” Ultrasound in Medicine and Biology, vol. 43, no. 4, pp. 790–803, 2017.

[45] G. A. Ateshian, C. A. Petersen, S. A. Maas, and J. A. Weiss, “A Numerical Scheme for Anisotropic Reactive Nonlinear Viscoelasticity,” Journal of Biomechanical Engineering, vol. 145, p. 011004, 08 2022.

[46] J. Bonet and R. D. Wood, Nonlinear Continuum Mechanics for Finite Element Analysis. Cambridge University Press, 2 ed., 2008.

[47] G. A. Holzapfel, T. C. Gasser, and R. W. Ogden, “A new constitutive framework for arterial wall mechanics and a comparative study of material models,” Journal of elasticity and the physical science of solids, vol. 61, pp. 1–48, 2000.

[48] A. Y. Malkin, “The use of a continuous relaxation spectrum for describing the viscoelastic properties of polymers,” Polymer Science Series A, vol. 48, pp. 39–45, Jan. 2006.

[49] S. A. Maas, B. J. Ellis, G. A. Ateshian, and J. A. Weiss, “Febio: Finite elements for biomechanics,” Journal of Biomechanical Engineering, vol. 134, p. 011005, 02 2012.

[50] B. M. Adams, W. J. Bohnhoff, K. R. Dalbey, M. S. Ebeida, J. P. Eddy, M. S. Eldred, R. W. Hooper, P. D. Hough, K. T. Hu, J. D. Jakeman, M. Khalil, K. A. Maupin, J. A. Monschke, E. M. Ridgway, A. A. Rushdi, D. T. Seidl, J. A. Stephens, and J. G. Winokur, *Dakota,* A Multilevel Parallel Object-Oriented Framework for Design Optimization, Parameter Estimation, Uncertainty Quantification, and Sensitivity Analysis: Version 6.15 User’s Manual, 11 2021.

[51] A. V. Tobolsky and T. D. Callinan, “Properties and structure of polymers,” Journal of The Electrochemical Society, vol. 107, p. 243C, oct 1960.

[52] R. Aspden, “The theory of fibre-reinforced composite materials applied to changes in the mechanical properties of the cervix during pregnancy,” Journal of Theoretical Biology, vol. 130, no. 2, pp. 213–221, 1988.

[53] C. J. Hansen, J. H. Rogers, A. J. Brown, N. Boatwright, S. Siricilla, C. M. O’Brien, S. Panja, C. M. Nichols, K. Devanathan, B. M. Hardy, M. D. Does, A. W. Anderson, B. C. Paria, A. Mahadevan-Jansen, J. Reese, and J. L. Herington, “Regional differences in three-dimensional fiber organization, smooth muscle cell phenotype, and contractility in the pregnant mouse cervix,” Science Advances, vol. 10, no. 51, p. eadr3530, 2024.

[54] X. Chen, D. Chen, E. Ban, K. C. Toussaint, P. A. Janmey, R. G. Wells, and V. B. Shenoy, “Glycosaminoglycans modulate long-range mechanical communication between cells in collagen networks,” Proceedings of the National Academy of Sciences, vol. 119, no. 15, p. e2116718119, 2022.

[55] Y. Akgul, R. A. Word, L. M. Ensign, Y. Yamaguchi, J. Lydon, J. Hanes, and M. Mahendroo, “Hyaluronan in cervical epithelia protects against infection-mediated preterm birth.,” The Journal of clinical investigation, vol. 124, pp. 5481–5489, Dec. 2014.

